# Edem1 activity in the fat body regulates insulin signalling and metabolic homeostasis in *Drosophila*

**DOI:** 10.1101/671305

**Authors:** Himani Pathak, Jishy Varghese

## Abstract

In *Drosophila,* nutrient status is sensed by the fat body, a functional homolog of mammalian liver and white adipocytes. The fat body conveys nutrient information to insulin-producing cells (IPCs) through humoral factors which regulate *Drosophila* insulin-like peptide (DILP) levels and insulin signalling. Insulin signalling has pleiotropic functions, which include the management of growth and metabolic pathways. Here, we report that Edem1 (<u>e</u>ndoplasmic reticulum <u>d</u>egradation-<u>e</u>nhancing α-<u>m</u>annosidase-like protein 1), an endoplasmic reticulum-resident protein involved in protein quality control, acts in the fat body to regulate insulin signalling and thereby the metabolic status in *Drosophila*. Edem1 limits the fat body derived *Drosophila* TNFα Eiger activity on IPCs and maintains systemic insulin signalling in fed conditions. During food deprivation, *edem1* gene expression levels drop, which aids in the reduction of systemic insulin signalling crucial for survival. Overall we demonstrate that Edem1 plays a vital role in helping the organism to endure a fluctuating nutrient environment by managing insulin signalling and metabolic homeostasis.

## Introduction

Energy homeostasis, the sum of all processes which maintain the balance between energy inflow and outflow; is vital for normal functioning, reproduction as well as longevity. Energy homeostasis in animals is brought about by the activity and interplay of various endocrine and neuroendocrine systems. Insulin/ Insulin-like growth factor (IGF) signalling pathway plays a significant role in the maintenance of energy balance and is well conserved in both vertebrates and invertebrates (Britton et al, 2002; Brogiolo et al, 2001; Clancy et al, 2001; Fabrizio et al, 2001; Fernandez & Torres-Aleman, 2012; Kenyon et al, 1993; Kimura et al, 1997). The perturbations in insulin signalling result in a plethora of effects like diabetes, obesity, reduced body size, resistance to starvation and oxidative stress (Accili et al, 1996; Bonafe et al, 2003; Britton et al., 2002; Clancy et al., 2001; Giannakou et al, 2004; Giannakou & Partridge, 2007; Holzenberger et al, 2003; Ikeya et al, 2002; Kahn et al, 2006; Katic & Kahn, 2005; Liu et al, 1993; Rulifson et al, 2002; Shimokawa et al, 2003; Sonntag et al, 2005; Tatar et al, 2001). *Drosophila melanogaster*, a widely used genetic model organism, has 8 insulin-like peptides (DILPs 1-8), which share structural and functional similarities with mammalian insulin and IGFs (Gronke et al, 2010a). Among these DILPs; DILP2, DILP3 and DILP5 are produced mainly by a subset of the median neurosecretory cells (mNSCs), the insulin-producing cells (IPCs), in the fly brain (Broughton et al, 2010; Geminard et al, 2009; Ikeya et al., 2002; Nassel, 2012). The major effector tissue of insulin signalling is the fat body, which is also the main energy reserve and nutrient sensor in flies (Geminard et al., 2009; Hwangbo et al, 2005). The fat body relays information about the nutrient status of the organism through humoral factors, which act on the IPCs directly or indirectly to control systemic insulin signalling (Agrawal et al, 2016; Bai et al, 2012; Colombani et al, 2003; Delanoue et al, 2016; Droujinine & Perrimon, 2016; Geminard et al., 2009; Ghosh & O’Connor, 2014; Koyama & Mirth, 2016; Rajan & Perrimon, 2013; Sano et al, 2015b; Sun et al, 2017). The fat derived signals that control IPC function include DILP6, a *Drosophila* insulin-like peptide; Unpaired2 (Upd2), a functional homolog of leptin in *Drosophila* and activator of JAK-STAT pathway; Eiger, the *Drosophila* Tumor Necrosis Factor α/ TNFα, which activates JNK signalling; CCHamide2, a nutrient responsive peptide hormone; Growth blocking peptide (GBP), a *Drosophila* cytokine; Stunted, a circulating insulinotropic peptide; female-specific independent of transformer (FIT) and activin-like ligand Dawdle. The molecular mechanisms that regulate the synthesis and secretion of the fat body derived signals (FDSs) is currently under intense investigation.

The Endoplasmic reticulum (ER) serves many functions in the eukaryotic cell, foremost of which is the synthesis and folding of nascent proteins with the help of molecular chaperones and folding enzymes. Hence, the ER is considered as the major quality-control site which ensures that only correctly folded proteins are allowed to leave to other cellular compartments. The ER is also considered to be the first storage site of secretory proteins and the ER activity is high in cells of endocrine and exocrine tissues due to the heavy protein trafficking in such cells. Genetic factors, physiological changes and fluctuations in the cellular environment might lead to misfolding of proteins (Liu & Kaufman, 2003) and the ER aids in eliminating proteins, which remain misfolded even after multiple rounds of folding attempts. Thus, a proper balance between the influx of proteins and the folding machinery in the ER is crucial for efficient protein quality control. When the ER homeostasis is upset misfolded proteins accumulate in the ER triggering an adaptive response called unfolded protein responses (UPR). The UPR signalling mainly involves three ER residing transmembrane sensors: inositol-requiring protein 1 (IRE1), activating transcription factor 6 (ATF6) and PKR-like ER kinase (PERK). The UPR sensors would initiate ER-associated degradation (ERAD) of terminally misfolded proteins, expand the ER membrane, increase the folding capacity of the ER and decrease the overall protein load in the ER (Liu & Kaufman, 2003). Permanently unfolded glycoproteins are recognised by ERAD-enhancing α-mannosidase-like proteins (Edem), which aid in the degradation of the misfolded proteins (Araki & Nagata, 2011; Kroeger et al, 2012; Molinari et al, 2003). Glycoproteins constitute a large proportion of proteins in a cell, hence the function of Edem is crucial for cellular homeostasis.

Here, we report that Edem1 activity in the *Drosophila* fat body is crucial for maintaining systemic insulin signalling. Down-regulation of *edem1* gene expression in the fat body resulted in the accumulation of DILP2 in the IPCs, a decrease of *dilp3* mRNA levels and reduced systemic insulin signalling, which led to nutrient imbalances and altered sensitivity to starvation. Our results also show that Edem1 regulates fat body derived *Drosophila* TNFα Eiger activity on the IPCs, crucial for managing systemic insulin signalling and metabolic status. Activation of target of rapamycin (TOR) signalling, the main amino acid sensor and a key regulator of Eiger activity rescued the effects of *edem1* down-regulation. Furthermore, in response to nutrient deprivation, *edem1* transcripts were found to be low, which we show is critical to the reduction of systemic insulin levels and better survival of flies during starvation. We propose that Edem1 acts as a key factor in the fat body, which maintains nutrient homeostasis by controlling the activity of the IPCs through Eiger.

## Results

### Edem1 maintains metabolic homeostasis

We embarked on a large scale genetic screen in *Drosophila* to identify factors that control nutrient homeostasis and insulin signalling. Towards this, we blocked various candidate genes, reported to be differentially expressed in the *miR-14* mutants that exhibited metabolic imbalances, in the *Drosophila* fat body using RNAi lines (Varghese et al, 2010a). We chose male flies for this study to minimize the effects of oogenesis on nutrient homeostasis. In this screen we identified Edem1, an ER-resident protein involved in protein quality control, as a putative regulator of metabolic status in *Drosophila*. Down-regulation of *edem1* transcripts in the fat body led to a significant increase in the levels of energy stores - triglycerides and glycogen, in 5 day old adult flies (Fig. 1A and B). In response to knock down of *edem1* in the fat body flies survived longer in response to acute nutrient deprivation (Fig. 1C). We chose five day old flies to completely avoid the influence of larval fat cells which persists in adult flies for few days after eclosion. We have confirmed the effects of blocking Edem1 in the fat body using independent RNAi lines, which rules out off-target effects and insertional site specific effects (Fig. S1B, C). The higher energy stores present in response to reduction of *edem1* levels in the fat body could account for the better survival of flies during nutrient deprivation (Fig. 1D and E). Along with changes in stored nutrient levels in adult flies, circulating glucose levels were high in the larval hemolymph (Fig. 1F). In addition, blocking *edem1* in the fat body led to enhanced feeding responses in the larvae (Fig. 1G), similar to responses reported earlier in food deprived larvae and also in response to low insulin due to its anorexigenic effects (Chouhan et al, 2017; Zhang et al, 2013). We also observed an increase in lifespan of the adult flies upon *edem1* down-regulation in the fat body (Fig. 1H). These data show that Edem1 function in the fat body is crucial in regulating metabolic homeostasis in *Drosophila*. The phenotypes observed in response to blocking *edem1* levels on larval circulating sugar levels, larval feeding, adult energy stores and life span indicated a reduction in insulin signalling, as similar phenotypes are reported in response to low insulin signalling (Bai et al, 2013; Bai et al., 2012; Bai et al, 2015; Bohni et al, 1999; Broughton et al, 2005; Clancy et al., 2001; Giannakou & Partridge, 2007; Gronke et al, 2010b; Haselton et al, 2010a; Hong et al, 2012; Hwangbo et al., 2005; Min et al, 2008; Partridge et al, 2011; Post et al, 2018; Shingleton et al, 2005; Slaidina et al, 2009; Tatar et al., 2001; Teleman et al, 2006; Varghese et al., 2010a; Wu et al, 2005). Next, we tested if insulin signalling is reduced in response to blocking *edem1* levels in the fat body.

**Fig 1.**
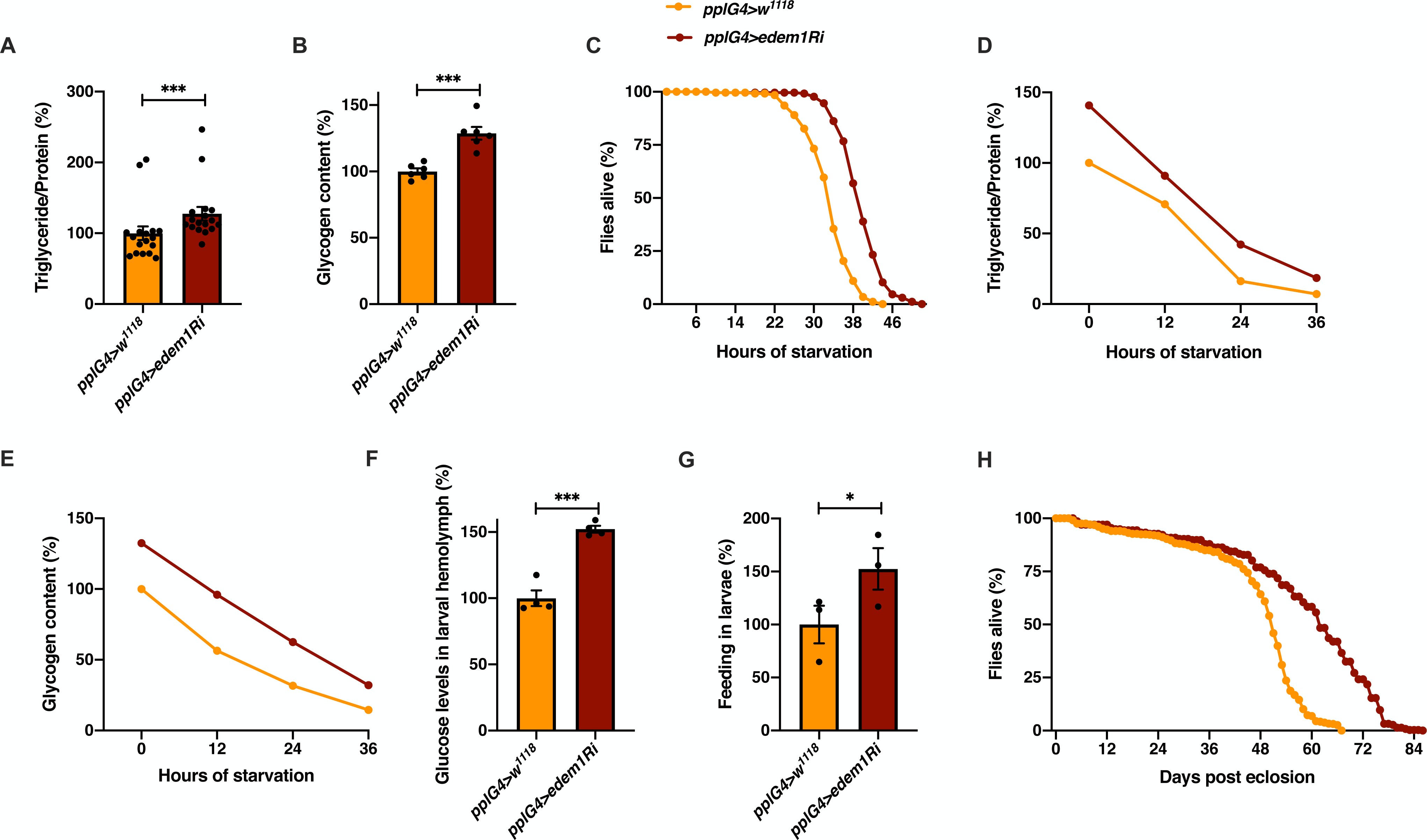
Edem1 maintains metabolic homeostasis. (A) Blocking *edem1* expression using RNAi in the fat body led to enhanced triglyceride levels in adult male flies. Data is shown as % ratio of triglyceride to total protein levels, normalised to 100% in *pplGal4>w^1118^* (control) and increase in experimental conditions *pplGal4>UAS-edem1-RNAi* [independent biological replicates = 17, P-value between control and *UAS-edem1-RNAi* is <0.001 (Mann-Whitney test)]. (B) Enhanced levels of glycogen in adult male flies caused by blocking *edem1* expression in the fat body. Data is shown as % of total glycogen levels, normalised to 100% in *pplGal4>w^1118^* (control) and increase in experimental conditions *pplGal4>UAS-edem1-RNAi* [independent biological replicates = 6, P-value between control and *UAS-edem1-RNAi* is <0.001 (Mann-Whitney test)]. (C) Enhanced resistance to starvation in adult male flies caused by blocking *edem1* expression in the fat body. Data shown as percentage of flies of *pplGal4>w^1118^* (control) and *pplGal4>UAS-edem1-RNAi* which were alive at various time points of starvation [independent biological replicates = 4, number of flies used for control is 255 and for *pplGal4>UAS-edem1-RNAi* is 262. P-value between control and *UAS-edem1-RNAi* is <0.001 (Log-rank test), Wald test = 189.8 on df = 1, p < 0.001 (cox-hazard proportional analysis)]. (D) and (E) Utilisation of triglycerides and glycogen at different stages of starvation upon *edem1* knockdown. Data is shown as % ratio of triglyceride to total protein levels in adult male flies, data is normalised to 100% in *pplGal4>w^1118^* (control) fed condition and change in response to indicated hours of starvation in control and experimental conditions *pplGal4>UAS-edem1-RNAi* is shown [independent biological replicates = 3, P-value between control and *UAS-edem1-RNAi* is 0.3844 (Log-rank test)]. Glycogen levels at different stages of starvation upon *edem1* knockdown. Data is normalised to 100% in *pplGal4>w^1118^* (control) fed condition and change in response to indicated hours of starvation in control and experimental conditions *pplGal4>UAS-edem1-RNAi* is shown [independent biological replicates = 3, P-value between control and *UAS-edem1-RNAi* is 0.0082 (Log-rank test)]. (F) Expression of *edem1* RNAi in the fat body led to enhanced glucose levels in the circulation. Data is shown as % of Glucose levels in the hemolymph, normalised to 100% in *pplGal4>w^1118^* (control) and increase in experimental conditions *pplGal4>UAS-edem1-RNAi* [independent biological replicates = 4, P-value between control and *UAS-edem1-RNAi* is <0.001 (Mann-Whitney test)]. (G) Blocking *edem1* gene expression in the fat body led to enhanced feeding responses in larvae. Data is shown as % food consumption in larvae, normalised to 100% in *pplGal4>w^1118^* (control) and increase in experimental conditions *pplGal4>UAS-edem1-RNAi* [independent biological replicates = 3, P-value between control and *UAS-edem1-RNAi* is 0.0329 (Welch’s t-test)]. (H) *edem1*-RNAi in the fat body led to enhanced life span in adult male flies. Data is shown as percentage of input flies *pplGal4>w^1118^, pplGal4>UAS-edem1-RNAi* which were alive across the days. [independent biological replicates = 3, number of flies used for control is 453 and for *pplGal4>UAS-edem1-RNAi* is 372. P-value between control and *UAS-edem1-RNAi* is <0.001 (Log-rank test), Wald test = 275.1 on df = 1, p < 0.001 (cox-hazard proportional analysis)]. **[*P-value *<0.05; ** <0.01,*** <0.001; Data information: In (A-B and F-G) data are presented as mean ± SEM*].**

### Edem1 function in the fat body maintains systemic insulin signalling

To measure the insulin signalling activity in response to blocking *edem1* in the fat body we checked gene expression of key downstream target genes of insulin pathway. Transcription of *4ebp (eIF4E-binding protein)*, *inr* (*insulin receptor*) and *dilp6* (*Drosophila insulin-like peptide 6*) is suppressed by insulin signalling and these insulin target genes can be used as a read out for insulin signalling activity (Puig et al, 2003; Slaidina et al., 2009). Blocking *edem1* in the fat body increased transcript levels of the insulin responsive genes, which indicate low insulin signalling (Fig. 2A). We speculated if Edem1 activity in the fat body could regulate IPC function and control systemic insulin signalling, as fat body is known to remotely control IPCs. To address whether Edem1 in the fat body regulates IPC function, the transcript levels of IPC specific DILPs - *dilp2, dilp3* and *dilp5* were measured in the late third instar larval stage. In response to the expression of *edem1*-RNAi in the fat body, *dilp3* mRNA levels were found to be low, however, there were no detectable changes in the mRNA levels of *dilp2* and *dilp5* (Fig. 2B). Previous studies report that nutrient deprivation would block DILP secretion from the IPCs into the hemolymph leading to an accumulation of DILPs and reduction in systemic insulin signalling (Geminard et al., 2009). We observed an increase of DILP2 puncta in IPCs in response to reducing *edem1* levels in the fat body when compared to that of control (Fig. 2C’ and C’’), which suggested an accumulation of DILP2 protein in the IPCs. Together, these observations suggest that *edem1* function in the fat body maintains systemic insulin signalling in the larvae by the regulation of IPC activity.

**Fig 2.**
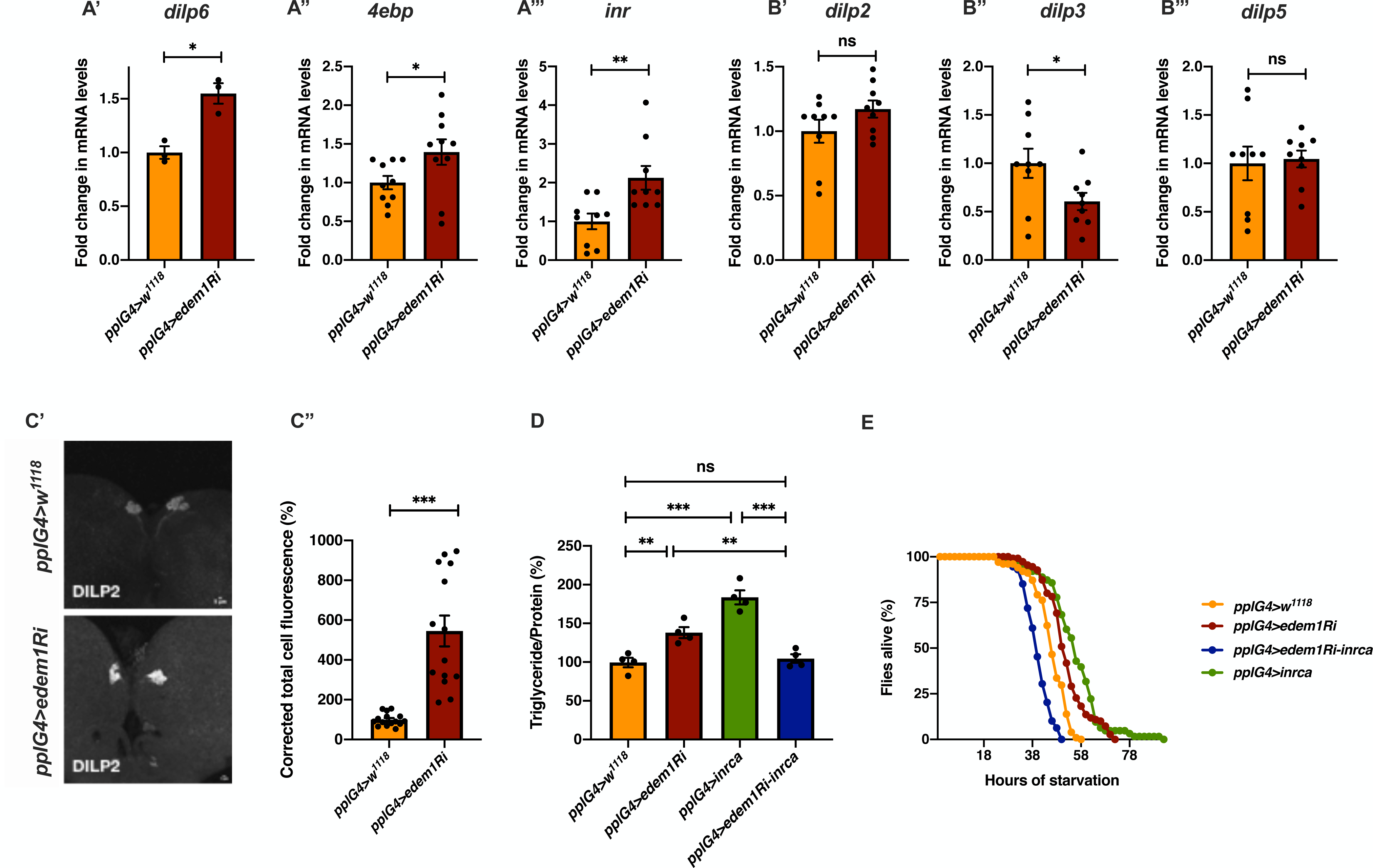
Blocking *edem1* in the fat body reduced insulin signalling. (A) Blocking *edem1* expression using RNAi in the fat body led to an increase of mRNA levels of insulin target genes *dilp6* (Fig. 2A’), *4ebp* (Fig. 2A’’) and *inr* (Fig. 2A’’’) in larvae. Data is shown as fold change in mRNA levels, values are normalised to *pplGal4>w^1118^* and fold change in *pplGal4>UAS-edem1-RNAi* is shown. [independent biological replicates = 3, P-value between control and *UAS-edem1-RNAi* is 0.0128 for *dilp6* (Welch’s t-test), for *4ebp* independent biological replicates = 10 and P-value is 0.0473 (Unpaired t test), for *inr* independent biological replicates = 9 and P-value is 0.0087 (Welch’s t test)]. (B) Blocking *edem1* expression using RNAi in the fat body also led to a decrease in the levels of IPC specific *dilp3* mRNA in larvae. Data is shown as fold change in mRNA levels, values are normalised to *pplGal4>w^1118^* and fold change in *pplGal4>UAS-edem1-RNAi* is shown. [n=9, P-value between control and *UAS-edem1-RNAi* is 0.1809 for *dilp2* (Mann-Whitney test), 0.0432 for *dilp3* (Welch’s t-test) and 0.8187 for *dilp5* (Welch’s t-test)]. (C) DILP2 protein in the larval IPCs shown as a representative image (Fig. 2C’) of anti-DILP2 antibody staining in larval brains of *pplGal4>w^1118^* [independent biological replicates = 15] and *pplGal4>UAS-edem1-RNAi* [independent biological replicates = 14]. Corrected total cell fluorescence values are normalised to *pplGal4>w^1118^* and fold change in *pplGal4>UAS-edem1-RNAi* is shown in Fig. 2C’’. [P-value between control and *UAS-edem1-RNAi* is <0.001 (Mann-Whitney test)] (D) Over expression of a constitutively active form of *inr* (*InR^A1325D^*) with *edem1* RNAi in the fat body led to the rescue of fat phenotype in adult male flies. Data is shown as % ratio of triglyceride to total protein levels, normalised to 100% in *pplGal4>w^1118^* (control) and changes in experimental conditions *pplGal4>UAS-edem1-RNAi*, *pplGal4>UAS-InR^A1325D^* and *pplGal4>UAS-edem1-RNAi; UAS-InR^A1325D^* [independent biological replicates = 4, P-value between control and *UAS-edem1-RNAi* is <0.001, P-value between control and *UAS*-*InR^A1325D^* is <0.001, P-value between *UAS-edem1-RNAi* and *UAS-edem1-RNAi*, *UAS*-*InR^A1325D^* is 0.001, P-value between *UAS-edem1-RNAi* and *UAS*-*InR^A1325D^* is 0.1292, P-value between *UAS*-*InR^A1325D^* and *UAS-edem1-RNAi*, *UAS*-*InR^A1325D^* is <0.001 and P-value between control and *UAS-edem1-RNAi*, *UAS*-*InR^A1325D^* is >0.9999 (Kruskal-Wallis test followed by Dunn’s post-hoc test)]. (E) Starvation resistance in adult male flies shown as percentage of input flies *pplGal4>w^1118^, pplGal4>UAS-edem1-RNAi, pplGal4>UAS*-*InR^A1325D^* and *pplGal4> UAS-edem1-RNAi*, *UAS*-*InR^A1325D^* which were alive at various time points of starvation [independent biological replicates = 3, number of flies used for control is 77, for *pplGal4>UAS-edem1-RNAi* is 110, for *pplGal4>UAS*-*InR^A1325D^* is 126 and for *pplGal4> UAS-edem1-RNAi*, *UAS*-*InR^A1325D^* is 128. P-value between control and *UAS-edem1-RNAi* is <0.001, P-value between control and *UAS*-*InR^A1325D^* is <0.001, P-value between *UAS-edem1-RNAi* and *UAS-edem1-RNAi*, *UAS*-*InR^A1325D^* is <0.001, P-value between *UAS-edem1-RNAi* and *UAS*-*InR^A1325D^* is 0.0038, P-value between *UAS*-*InR^A1325D^* and *UAS-edem1-RNAi*, *UAS*-*InR^A1325D^* is <0.001 and P-value between control and *UAS-edem1-RNAi*, *UAS*-*InR^A1325D^* is <0.001 (Log-rank test), Wald test = 10.53 on df = 1, p=0.001 (cox-proportional hazard analysis)]. **[*P-value *<0.05; ** <0.01,*** <0.001; Data information: In (A-B, C’’ and D) data are presented as mean ± SEM*].**

As the next approach we tested whether the reduction of insulin signalling in response to blocking *edem1* in the fat body was responsible for the metabolic phenotypes. A constitutively active form of insulin receptor (*InR^A1325D^*) was co-expressed with *edem1*-RNAi in the fat body. *InR^A1325D^,* which harbours an Ala-Asp mutation at position 1325, would activate downstream insulin signalling independent of DILP ligand and hence should alleviate phenotypes caused by low insulin signalling (Broughton et al., 2005; DiAngelo & Birnbaum, 2009; Kannan & Fridell, 2013; Tettweiler et al, 2005b). As expected, expression of *InR^A1325D^* was sufficient to alleviate high triglyceride levels and starvation resistance observed in response to knock down of *edem1* levels in the fat body (Fig. 2D and E). We have performed experiments using UAS-*control* transgenes to rule out the effect of Gal4 titration in this experiment and all future rescue experiments where multiple UAS transgenes are driven by the same Gal4 driver (data not shown). Thus, blocking *edem1* in the fat body reduced systemic insulin signalling, which led to metabolic phenotypes. These experiments confirm that Edem1 function in the fat body is crucial to maintain systemic insulin signalling and metabolic homeostasis.

### Fat body derived signals are involved in Edem1 mediated regulation of IPCs

*Drosophila* fat body controls IPC function with the aid of a set of humoral factors, which relays the nutritional status of the organism to the IPCs. The fat body derived signals (FDSs) control DILP release from the IPCs into the hemolymph leading to effects on growth and maintenance of metabolic balance. In addition, changes in *dilp* gene expression has also been reported in response to fat body derived signals. We next investigated whether blocking Edem1 led to changes in the levels of FDSs and thereby the function of IPCs.

To test the role of FDSs in *edem1* knock down phenotypes we measured the levels or activity of various FDSs. We saw an increase in the transcript levels of *dilp6* in response to knock down of *edem1* in the fat body (Fig. 2A’). Next, we measured the mRNA levels of *upd2*, *totA* and the levels of STAT92E-GFP as read outs for activity of JAK-STAT pathway, a cell signalling pathway activated by Upd2, an FDS reported to regulate IPC functions (Rajan & Perrimon, 2013). Blocking *edem1* expression in the fat body led to a decrease in *totA* and *upd2* mRNA levels (Fig. S2A’ and A’’). In addition, STAT92E-GFP expression in the brain was found to be low in response to reduced *edem1* in the fat body (Fig. S2B’ and B’’). We also measured Drosophila TNFα Eiger levels, another FDS that acts on IPCs through it’s receptor Grindelwald, and activation of downstream JNK signalling by Eiger. The transcript levels of tace, the <u>T</u>NF<u>α</u> <u>c</u>onverting <u>e</u>nzyme encoding gene; *eiger* and *Nlaz*, a key target of JNK signalling were measured (Agrawal et al., 2016; Hull-Thompson et al, 2009; Pasco & Leopold, 2012), and the levels of these genes were found to be increased in response to *edem1*-RNAi (Fig. 3A, B and D). The cleaved form of Eiger protein in the whole body extracts was also found to be higher in *edem1*-RNAi (Fig. 3C’ and C’’), which confirmed enhanced levels of active form of Eiger released by blocking Edem1 function in the fat body. Eiger and Dilp6 are considered to be negative regulators of IPC function, whereas, Upd2 is expected to activate IPCs (Agrawal et al., 2016; Bai et al., 2012; Rajan & Perrimon, 2013), the gene expression changes observed here suggested that these FDSs might mediate the effects of *edem1* knock down on insulin signalling.

**Fig 3.**
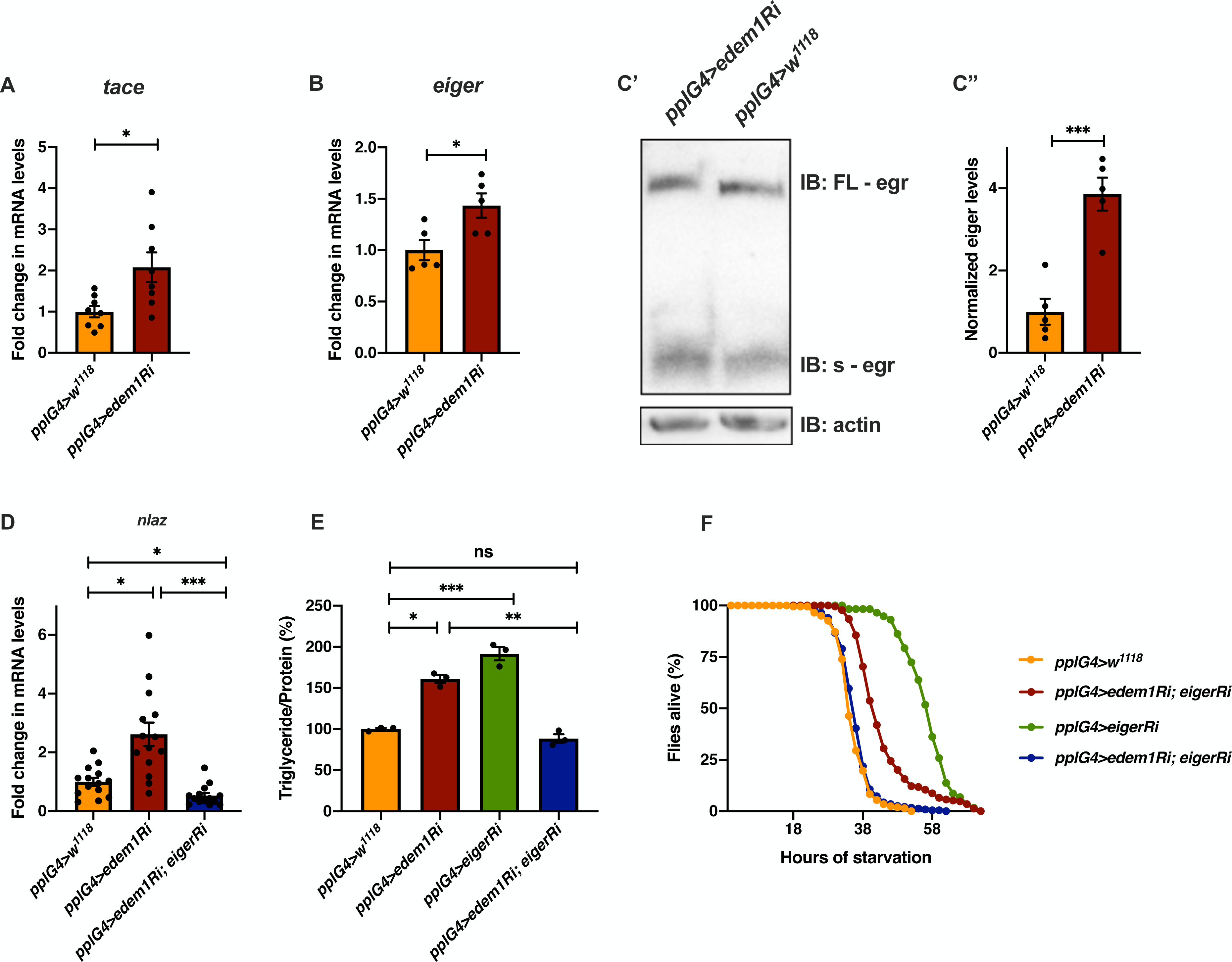
Reduction of *edem1* levels in the fat body enhanced JNK signalling. (A) Blocking *edem1* expression using RNAi in the fat body led to an increase of mRNA levels of *tace*. Data is shown as fold change in mRNA levels, values are normalised to *pplGal4>w^1118^* and fold change in *pplGal4>UAS-edem1-RNAi* is shown. [independent biological replicates = 8, P-value between control and *UAS-edem1-RNAi* is 0.0211 (Welch’s t-test)]. (B) Blocking *edem1* expression using RNAi in the fat body led to an increase of mRNA levels of *eiger*. Data is shown as fold change in mRNA levels, values are normalised to *pplGal4>w^1118^* and fold change in *pplGal4>UAS-edem1-RNAi* is shown. [independent biological replicates = 5, P-value between control and *UAS-edem1-RNAi* is 0.0226 (Welch’s t-test)]. (C) Blocking *edem1* expression using RNAi in the fat body led to an increase in the circulating levels of *eiger* (Fig. 3C’). Data is shown as fold change in normalized soluble eiger levels (Fig. 3C’’), values are normalised to *pplGal4>w^1118^* and fold change in *pplGal4>UAS-edem1-RNAi* is shown. [independent biological replicates = 5, P-value between control and *UAS-edem1-RNAi* is 0.0006 (Welch’s t-test)]. (D) Expression of *eiger* RNAi in the fat body rescued increase in *NLaz* mRNA levels caused by blocking *edem1* levels in the fat body, data is shown as fold change in mRNA levels. Data is shown as fold change in mRNA levels, values are normalised to *pplGal4>w^1118^* and fold change in *pplGal4>UAS-edem1-RNAi, pplGal4>UAS-eiger-RNAi* and *pplGal4>UAS-edem1-RNAi; UAS-eiger-RNAi* is shown. [independent biological replicates = 14. P-value between control and *UAS-edem1-RNAi* is 0.0171, P-value between *UAS-edem1-RNAi* and *UAS-edem1-RNAi; UAS-eiger-RNAi* is <0.001 and between control and *UAS-edem1-RNAi; UAS-eiger-RNAi* is 0.1085 (Kruskal-Wallis test followed by Dunn’s post-hoc test)]. (E) Expression of *eiger* RNAi in the fat body rescued increase in triglyceride caused by blocking *edem1* levels in the fat body. Data is shown as % ratio of triglyceride to total protein levels, normalised to 100% in *pplGal4>w^1118^* (control) and changes in experimental conditions *pplGal4>UAS-edem1-RNAi* and *pplGal4>UAS-edem1-RNAi; UAS-eiger-RNAi* is shown. [independent biological replicates = 3, P-value between control and *UAS-edem1-RNAi* is 0.0052, P-value between control and *UAS-eiger-RNAi* is <0.001, P-value between *UAS-edem1-RNAi* and *UAS-edem1-RNAi*, *UAS-eiger-RNAi* is <0.001, P-value between *UAS-edem1-RNAi* and *UAS-eiger-RNAi* is 0.8315, P-value between *UAS-eiger-RNAi* and *UAS-edem1-RNAi*, *UAS-eiger-RNAi* is <0.001 and P-value between control and *UAS-edem1-RNAi*, *UAS-eiger-RNAi* is >0.9999 (Kruskal-Wallis test followed by Dunn’s post-hoc test)]. (F) Enhanced starvation resistance shown as percentage of flies which were alive at various time points of starvation in the following genotypes - *pplGal4>w^1118^*, *pplGal4>UAS-edem1-RNAi* and *pplGal4> UAS-edem1-RNAi; UAS-eiger-RNAi* is shown. [independent biological replicates = 3, number of flies used for control is 203, for *pplGal4>UAS-edem1-RNAi* is 229, for *pplGal4>UAS*-*eiger-RNAi* is 58 and for *pplGal4> UAS-edem1-RNAi*, *UAS*-*Ieiger-RNAi* is 233. P-value between control and *UAS-edem1-RNAi* is <0.001, P-value between control and *UAS-eiger-RNAi* is <0.001, P-value between *UAS-edem1-RNAi* and *UAS-edem1-RNAi*, *UAS-eiger-RNAi* is <0.001, P-value between *UAS-edem1-RNAi* and *UAS-eiger-RNAi* is <0.001, P-value between *UAS-eiger-RNAi* and *UAS-edem1-RNAi*, *UAS-eiger-RNAi* is <0.001 and P-value between control and *UAS-edem1-RNAi*, *UAS-eiger-RNAi* is 0.0459 (Log-rank test), Wald test = 0.09 on df = 1, p=0.8 (cox-proportional hazard analysis)]. **[*P-value *<0.05; ** <0.01,*** <0.001; Data information: In (A-B, C’’-E) data are presented as mean ± SEM*].**

### Eiger is involved in Edem1 mediated regulation of IPCs

To identify the FDS(s) involved in mediating the metabolic phenotypes observed by blocking *edem1* levels, we expressed *eiger-*RNAi, *dilp6-*RNAi or *upd2*, together with *edem1*-RNAi in the fat body. Down regulation of *eiger* mRNA and over expression of *upd2* rescued *edem1* knock down phenotypes of lipid stores and starvation resistance, however, there were no effects with *dilp6*-RNAi (Fig. 3E and F, Fig. S2C-F). This suggested that either Upd2 or Eiger could be regulated by Edem1 and manage IPC function. We did not see any significant changes to DILP2 levels in the IPCs, *dilp3* mRNA levels and insulin target genes by overexpression of *upd2* in fat body that express *edem1*-RNAi (data not shown). However, when we down-regulated *eiger* mRNA in *edem1*-RNAi expressing fat body, we observed a reduction of DILP2 puncta in the IPCs seen in response to *edem1*-RNAi expression in the fat body (Fig. 4A’ and A’’). In addition, transcript levels of *dilp3* (Fig. 4B) and insulin target genes *dilp6* (Fig. 4C’) *and inr* (Fig. 4C’’) were restored by reducing Eiger levels in the *edem1*-RNAi back ground. Furthermore, the increase of glucose levels in the hemolymph seen in response to blocking *edem1* was suppressed by co-expression of *eiger*-RNAi (Fig. 4D). Thus, Edem1 mediated regulation of Eiger is crucial for managing insulin levels and nutrient homeostasis, whereas the Edem1 mediated regulation of Upd2 function manages nutrient homeostasis but doesn’t do so by acting at the level of insulin signalling.

**Fig 4.**
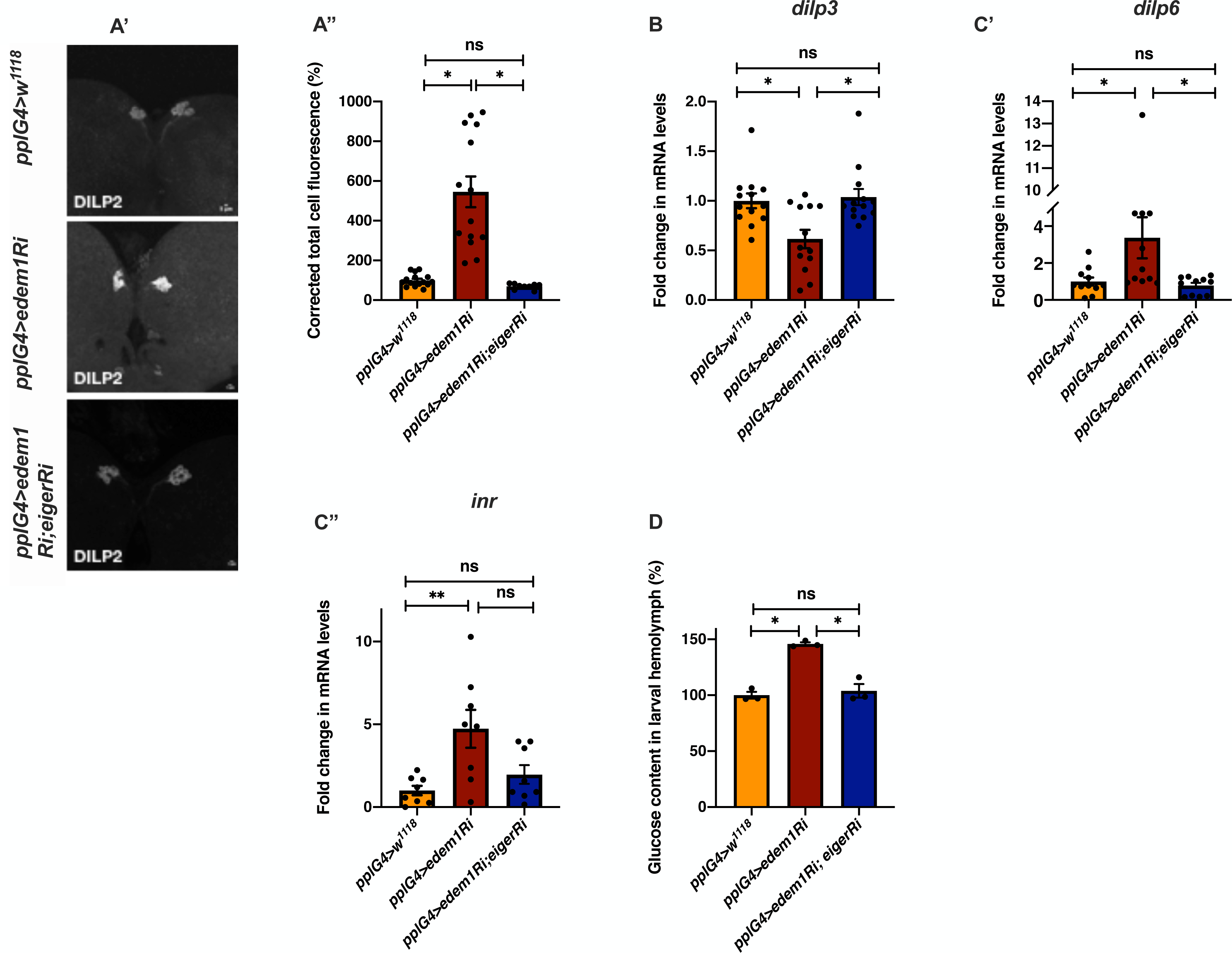
Knock down of *eiger* in the fat body rescued the metabolic phenotypes. (A) DILP2 levels in the IPCs in response to *edem1*-RNAi was rescued by reducing *eiger* in the fat body (Fig. 4A’). Shown are representative images of anti-DILP2 antibody staining in larval brains of *pplGal4>w^1118^* [independent biological replicates = 15]; *pplGal4>UAS-edem1-RNAi* [independent biological replicates = 14] and *pplGal4> UAS-edem1-RNAi; UAS-eiger-RNAi* [independent biological replicates = 9]. Corrected total cell fluorescence values are normalised to *pplGal4>w^1118^* and fold change in *pplGal4>UAS-edem1-RNAi* and *pplGal4> UAS-edem1-RNAi; UAS-eiger-RNAi* is shown in Fig. 4A’’. [P-value between control and *UAS-edem1-RNAi* is <0.001, P-value between control and *UAS-edem1-RNAi; UAS-eiger-RNAi* is >0.9999 and P-value between *UAS-edem1-RNAi* and *UAS-edem1-RNAi; UAS-eiger-RNAi* is <0.001 (Kruskal-Wallis test followed by Dunn’s post-hoc test)] (B) Reduction of *dilp3* mRNA levels, in response to *edem1*-RNAi was rescued by reducing *eiger* in the fat body, data is shown as fold change in mRNA levels, values are normalised to *pplGal4>w^1118^* fold change in *pplGal4>UAS-edem1-RNAi* and *pplGal4> UAS-edem1-RNAi; UAS-eiger-RNAi* is shown [independent biological replicates = 13. P-value between control and *UAS-edem1-RNAi* is 0.0395, between control and *UAS-edem1-RNAi; UAS-eiger-RNAi* is >0.9999 and between *UAS-edem1-RNAi* and *UAS-edem1-RNAi; UAS-eiger-RNAi* is 0.0253 (Kruskal-Wallis test followed by Dunn’s post-hoc test)]. (C) Increase in *dilp6,* in response to *edem1*-RNAi was rescued by reducing *eiger* in the fat body (Fig. 4C’) and *inr* (Fig. 4C’’). Data is shown as fold change in mRNA levels, values are normalised to *pplGal4>w^1118^* and fold change in *pplGal4>UAS-edem1-RNAi* and *pplGal4> UAS-edem1-RNAi; UAS-eiger-RNAi* is shown [independent biological replicates = 11 for *dilp6* and independent biological replicates = 8 *for inr*. P-value between control and *UAS-edem1-RNAi* is 0.0392 for *dilp6* and 0.0194 for *inr*. P-value between control and *UAS-edem1-RNAi; UAS-eiger-RNAi* is >0.9999 *for dilp6* and >0.9999 for *inr*, P-value between *UAS-edem1-RNAi* and *UAS-edem1-RNAi; UAS-eiger-RNAi* is 0.0296 for *dilp6* and 0.1976 for *inr* (Kruskal-Wallis test followed by Dunn’s post-hoc test)]. (D) Increase in Glucose levels in the hemolymph in response to *edem1*-RNAi was rescued by reducing *eiger* in the fat body. Data is shown as % of Glucose levels in the hemolymph, normalised to 100% in *pplGal4>w^1118^* (control) and changes in experimental conditions *pplGal4>UAS-edem1-RNAi* and *pplGal4> UAS-edem1-RNAi; UAS-eiger-RNAi is shown* [independent biological replicates = 3, P-value between control and *UAS-edem1-RNAi* is 0.0105, P-value between control and *UAS-edem1-RNAi; UAS-eiger-RNAi* is >0.9999, P-value between *edem1-RNAi* and *UAS-edem1-RNAi; UAS-eiger-RNAi* is 0.0175 (Kruskal-Wallis test followed by Dunn’s post-hoc test)]. **[*P-value *<0.05; ** <0.01,*** <0.001; Data information: In (A’’-D) data are presented as mean ± SEM*].**

The fat body derived cytokine Eiger is an upstream activator of c-Jun N-terminal kinase (JNK) pathway in flies and previous studies have shown that JNK signalling extends life span and limits growth by antagonizing cellular and organism-wide responses to insulin signalling (Agrawal et al., 2016; Andersen et al, 2015; Hirosumi et al, 2002; Oh et al, 2005; Wang et al, 2005). The increase of *NLaz* transcript levels in response to blocking *edem1* levels in the fat body was rescued completely by the co-expression of *eiger*-RNAi, showing that the increase of JNK signalling in response to blocking *edem1* expression in the fat body is due to enhanced Eiger levels (Fig. 3D). These experiments confirm that Edem1 function in the fat body regulates Eiger activity and JNK signalling.

Next, we carried out experiments to confirm if regulation of Eiger by Edem1 is crucial for maintaining IPC function and metabolic homeostasis. Towards this we used two approaches: (i) we activated TOR signalling pathway in *edem1*-RNAi expressing fat body (Fig. 5), as TOR signalling has been shown to block Eiger activation and (ii) we performed co-culture experiments by blocking the TNF receptor *grindelwald* in the IPCs (Fig. 6). TOR signalling pathway regulates a plethora of cellular processes including cell growth, proliferation cell survival, etc., depending on nutrient levels. Recently TOR has been reported to repress *tace* transcription, which would in turn suppress the production of active Eiger from the fat body (Agrawal et al., 2016). Rheb (Ras homolog enriched in brain), a member of Ras superfamily of GTP binding proteins, activates TOR kinase and results in growth and regulation of metabolic pathways (Garami et al, 2003; Oldham, 2011; Oldham et al, 2000; Saucedo, 2003). Increase in the levels of JNK pathway target *NLaz* in response to *edem1* downregulation in the fat body was abrogated by over-expression of *rheb* (Fig. 5A), which confirms that activating TOR signalling is sufficient to suppress JNK signalling possibly by the regulation of Eiger activity. Changes in the reduction of insulin signalling (Fig. 5B), *dilp3* transcript levels (Fig. 5C), excess fat levels (Fig. 5D) and starvation resistance (Fig. 5E) in response to blocking Edem1 levels were rescued by the over expression of *rheb* in the fat body. This confirmed that Edem1 mediated regulation of JNK activity is crucial for managing systemic insulin signalling, fat storage and starvation sensitivity.

**Fig 5.**
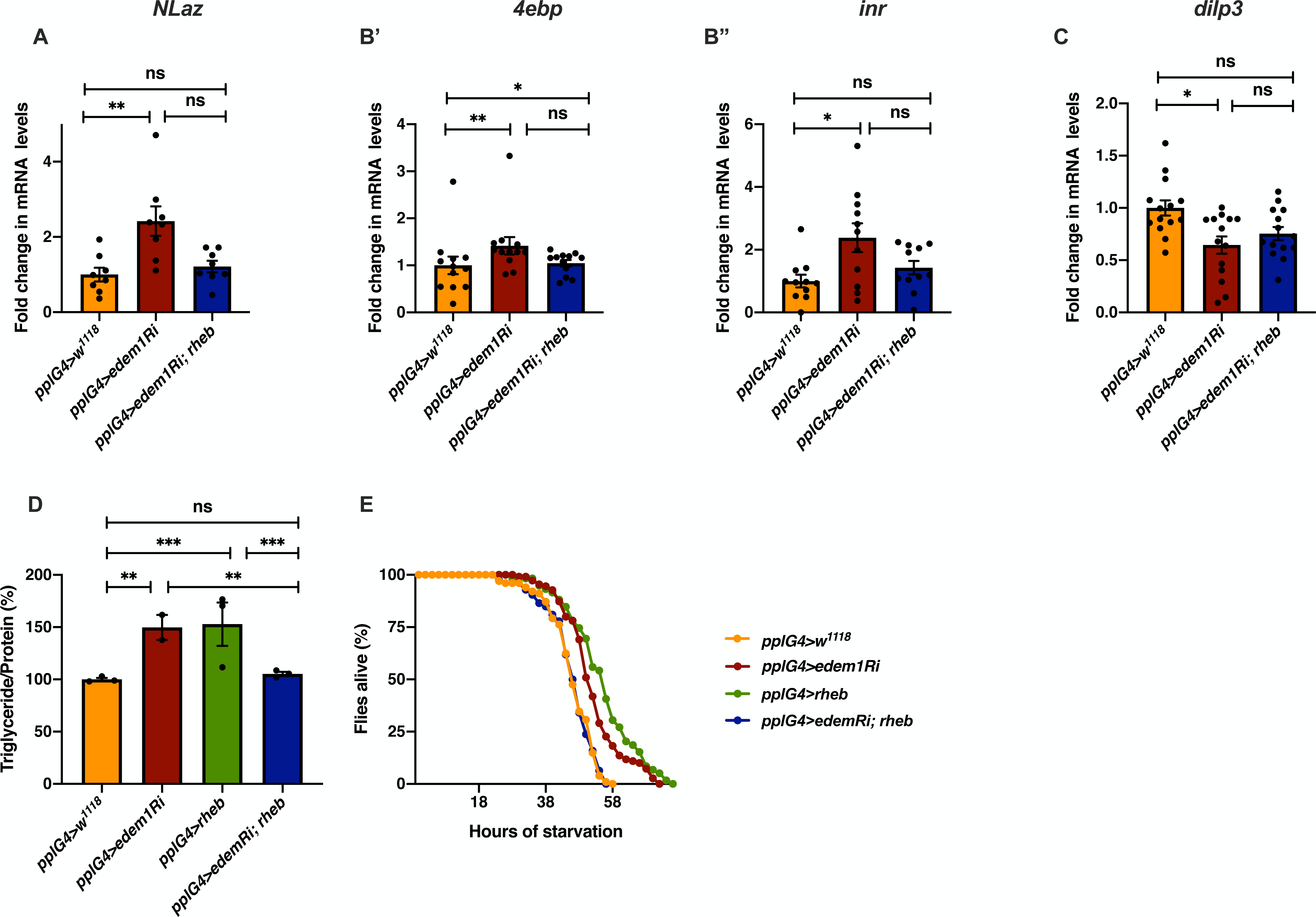
Activating TOR signalling rescued *edem1* mediated phenotypes. (A) Increase in mRNA levels of *NLaz* in response to blocking *edem1* expression was rescued by co-expression of *UAS-rheb*. Data is shown as fold change in mRNA levels, values are normalised to *pplGal4>w^1118^* and fold change in *pplGal4>UAS-edem1-RNAi* and *pplGal4> UAS-edem1-RNAi; UAS-rheb* is shown. [independent biological replicates = 8. P-value between control and *UAS-edem1-RNAi* is 0.0039, P-value between *UAS-edem1-RNAi* and *UAS-edem1-RNAi; UAS-rheb* is 0.0486, P-value between control and *UAS-edem1-RNAi; UAS-rheb* is >0.9999 (Kruskal-Wallis test followed by Dunn’s post-hoc test)]. (B) Increase of mRNA levels of *4ebp* (Fig. 5B’) and *inr* (Fig. 5B’’) in response to blocking *edem1* expression was alleviated by co-expression of *UAS-rheb*. Data is shown as fold change in mRNA levels, values are normalised to *pplGal4>w^1118^* and fold change in *pplGal4>UAS-edem1-RNAi* and *pplGal4> UAS-edem1-RNAi; UAS-rheb* is shown. [independent biological replicates = 12 for *4ebp*. P-value between control and *edem1*-RNAi is 0.0085, P-value between *UAS-edem1-RNAi* and *UAS-edem1-RNAi; UAS-rheb* is 0.1143, P-value between control and *UAS-edem1-RNAi; UAS-rheb* is >0.9999 (Kruskal-Wallis test followed by Dunn’s post-hoc test); independent biological replicates = 11 for *inr*. P-value between control and *UAS-edem1-RNAi* is 0.0406, P-value between *UAS-edem1-RNAi* and *UAS-edem1-RNAi; UAS-rheb* is 0.7277, P-value between control and *UAS-edem1-RNAi; UAS-rheb* is 0.5799 (Kruskal-Wallis test followed by Dunn’s post-hoc test)]. (C) Decrease in mRNA levels of *dilp3* in response to blocking *edem1* expression was rescued by co-expression of *UAS-rheb*. Data is shown as fold change in mRNA levels, values are normalised to *pplGal4>w^1118^* and fold change in *pplGal4>UAS-edem1-RNAi* and *pplGal4> UAS-edem1-RNAi; UAS-rheb* is shown. [independent biological replicates = 14. P-value between control and *UAS-edem1-RNAi* is 0.0231, P-value between *UAS-edem1-RNAi* and *UAS-edem1-RNAi; UAS-rheb* is >0.9999, P-value between control and *UAS-edem1-RNAi; UAS-rheb* is 0.1356 (Kruskal-Wallis test followed by Dunn’s post-hoc test)]. (D) Over-expression of *rheb* in the fat body rescued enhanced stored fat levels. Data is shown as % ratio of triglyceride to total protein levels, values are normalised to *pplGal4>w^1118^* and fold change in *pplGal4>UAS-edem1-RNAi* and *pplGal4> UAS-edem1-RNAi; UAS-rheb* is shown. [independent biological replicates = 4, P-value between control and *UAS-edem1-RNAi* is 0.0003, P-value between control and *UAS-rheb* is <0.001, P-value between *UAS-edem1-RNAi* and *UAS-edem1-RNAi*, *UAS-rheb* is 0.0005, P-value between *UAS-edem1-RNAi* and *UAS-rheb* is 0.5184, P-value between *UAS-rheb* and *UAS-edem1-RNAi*, *UAS-rheb* is <0.001 and P-value between control and *UAS-edem1-RNAi*, *UAS-rheb* is >0.9999 (Kruskal-Wallis test followed by Dunn’s post-hoc test)]. (E) Over-expression of *rheb* in the fat body rescued increased starvation resistance. Data is shown as percentage of flies which were alive at various time points of starvation in the following genotypes - *pplGal4>w^1118^*, *pplGal4>UAS-edem1-RNAi, pplGal4>UAS-rheb* and *pplGal4> UAS-edem1-RNAi; UAS-rheb.* [independent biological replicates = 5, number of flies used for control is 101, for *pplGal4>UAS-edem1-RNAi* is 110, for *pplGal4>UAS*-*rheb* is 59 and for *pplGal4> UAS-edem1-RNAi*, *UAS*-*rheb* is 126. P-value between control and *UAS-edem1-RNAi* is <0.0001, P-value between control and *UAS-edem1-RNAi* is <0.001, P-value between control and *UAS-rheb* is <0.001, P-value between *UAS-edem1-RNAi* and *UAS-edem1-RNAi*, *UAS-rheb* is <0.001, P-value between *UAS-edem1-RNAi* and *UAS-rheb* is 0.0247, P-value between *UAS-rheb* and *UAS-edem1-RNAi*, *UAS-rheb* is <0.001 and P-value between control and *UAS-edem1-RNAi*, *UAS-rheb* is 0.9628 (Log-rank test), Wald test = 0.21 on df = 1, p=0.6 (cox-proportional hazard analysis)]. **[*P-value *<0.05; ** <0.01,*** <0.001; Data information: In (A-D) data are presented as mean ± SEM*].**

**Fig. 6.**
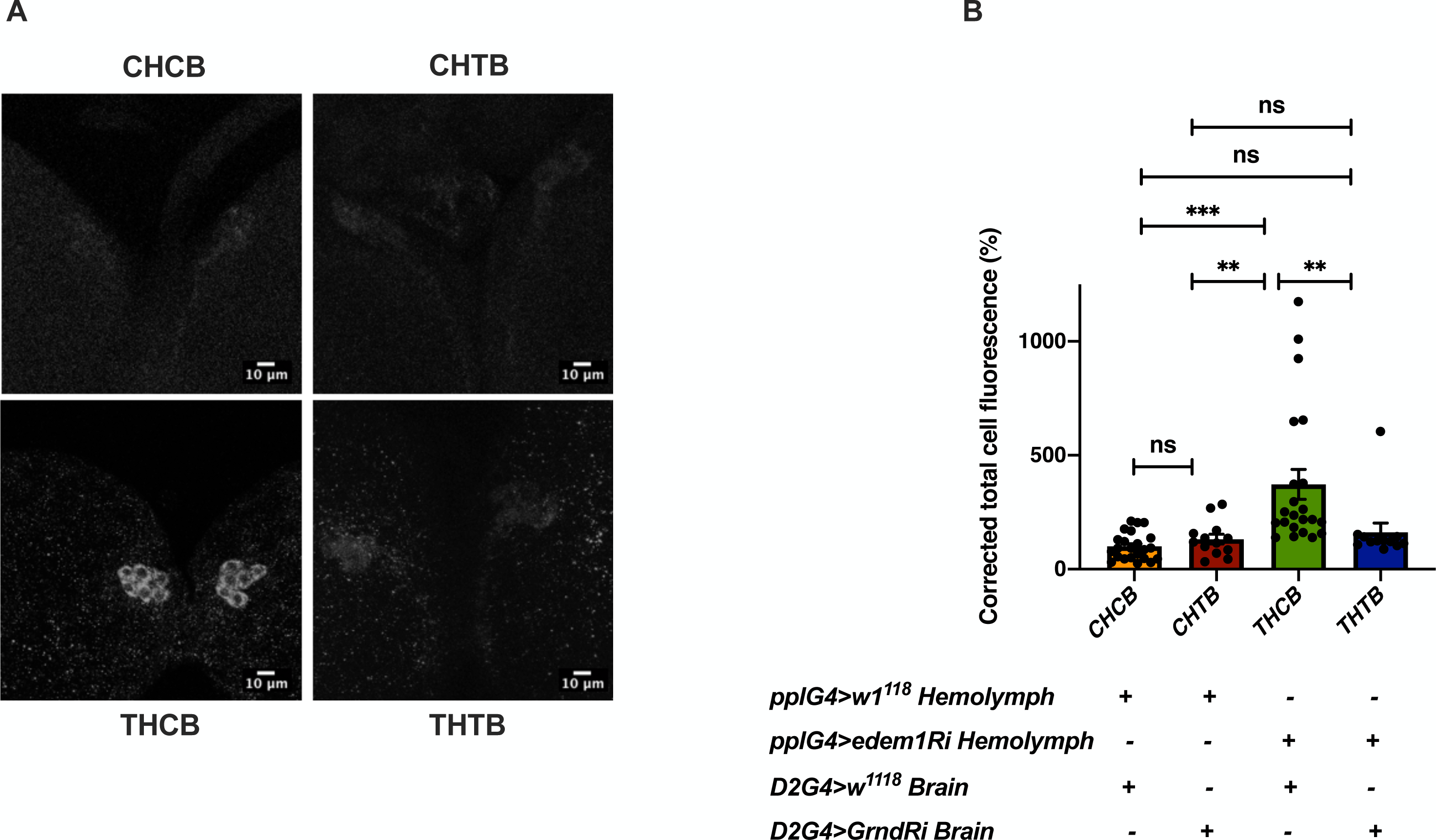
Blocking *grindelwald* in the IPCs rescued the increased accumulation of DILP2. (A) DILP2 accumulation in the IPCs in response to *edem1*-RNAi was rescued by blocking Eiger receptor *grnd* in the IPCs. Representative images of anti-DILP2 antibody staining in larval brains of *Dilp2Gal4>w^1118^* treated with hemolymph from *pplGal4*>*w^1118^* [independent biological replicates = 23] (CHCB), *Dilp2Gal4>UAS-grnd-RNAi* treated with hemolymph from *pplGal4*>*w^1118^* [independent biological replicates = 13] (CHTB), *Dilp2Gal4>w^1118^* incubated with hemolymph from *pplGal4>UAS-edem1-RNAi* [independent biological replicates = 22] (THCB) and *Dilp2Gal4>UAS-grnd-RNAi* incubated with hemolymph from *pplGal4>UAS-edem1-RNAi* [independent biological replicates = 12] (THTB) larvae are shown. (B) Corrected total cell fluorescence values are normalised to CHCB and fold change in CHTB, THCB and THTB are shown. [P-value between CHCB and CHTB is >0.9999, P-value between CHCB and THCB is <0.001, P-value between CHCB and THTB is >0.9999, P-value between CHTB and THCB is 0.0018, P-value between CHTB and THTB is >0.9999 and P-value between THCB and THTB is 0.0085 (Kruskal-Wallis test followed by Dunn’s post-hoc test)]. **[*P-value *<0.05; ** <0.01,*** <0.001; Data information: In (B) data are presented as mean ± SEM*].**

Next, we tried to show that Edem1 function in the fat body regulates Eiger activity on IPCs. In low protein diet, the soluble form of Eiger binds to its receptor Grindelwald (Grnd) in the IPCs, thereby activate JNK signalling and suppress *dilp* transcript levels (Agrawal et al., 2016; Andersen et al., 2015). We performed *ex vivo* organ culture experiments, using hemolymph isolated from control larvae and larvae in which *edem1* expression in the fat body was blocked. Brains dissected from wild type larvae and larvae with *grnd* knock down in the IPCs, were incubated with the hemolymph from the above mentioned conditions. As expected, treatment of control larval brains with hemolymph from *edem1*-RNAi larvae (THCB) led to accumulation of DILP2 in the IPCs, when compared to hemolymph from control larvae (CHCB) (Fig. 6A and B). We observed less DILP2 puncta in IPCs in response to blocking *grnd* (THTB) when compared to control IPCs treated with hemolymph from *edem1*-RNAi larvae (THCB) (Fig. 6A and B). Blocking *grnd* did not change the levels of DILP2 in the IPCs as *IPC>grnd-RNAi* brains did not show any changes in DILP2 puncta when treated with hemolymph extracted from control larvae (CHTB). Thus, accumulation of DILP2 in the IPCs in response to blocking *edem1* levels in the larval fat body is due to Eiger activity on the IPCs through TNF receptor Grindelwald. Together these results confirm that Edem1 activity in the fat body regulates Eiger mediated JNK signalling in the IPCs and manage systemic insulin signalling and metabolic status of flies.

### Reduction in *edem1* levels during starvation is crucial for survival

Insulin signalling aids an organism to respond to changes in the nutrient environment by managing various biological functions. In response to nutrient deprivation insulin signalling is reduced, which would allow an organism to manage its energy stores and induce various hunger triggered behavioral responses (Arsic & Guerin, 2008; Britton et al., 2002; Erion & Sehgal, 2013). Our experiments show that lowering *edem1* levels improved survival against starvation by reducing insulin signalling (Fig. 1C and Fig. 2E). Hence, we tested if *edem1* levels are lowered during nutrient deprivation, which may aid in better survival of flies by the reduction of insulin levels. Moreover, blocking *edem1* in the fat body in larvae was sufficient to enhance the appetite, similar to hunger induced responses observed in food deprived larvae (Fig. 1G). As expected, in response to food depletion we observed a reduction of *edem1* mRNA levels (Fig. 7A). To test whether reduction of *edem1* levels is essential for protection against starvation we over expressed *edem1* in the fat body in food deprived flies and checked their starvation responses. Overexpression of *edem1* in the fat body reduced the survival of flies during food deprivation (Fig. 7B’, Fig. S3A), confirming that the reduction of *edem1* levels is crucial for survival in response to nutrient depletion. Enhancing the levels of Edem1 during starvation affected mobilisation of stored fat (Fig 7B’’). Higher Edem1 levels during starvation blocked the increase of *eiger* transcripts (Fig 7C) although the levels of the cleaved form of Eiger did not get affected either in fed or starved conditions (data not shown). In response to acute starvation JNK signalling is increased and the transcript levels of *dilp3* and insulin signalling are decreased (Fig. 7D - F). These effects of starvation was abrogated by *edem1* over- expression in the fat body (Fig. 7C - F). Meanwhile, over-expression of Edem1 in the fed conditions did not affect fat levels and insulin signalling, except for a slight change in *dilp6* levels (7C - F). Thus, elevated Edem1 expression affects metabolic homeostasis and survival during starvation conditions. Here, we conclude that lowering of *edem1* transcripts in the fat body during starvation facilitates activation of Eiger and reduction of insulin signalling, which results in the enhanced survival of flies.

**Fig. 7.**
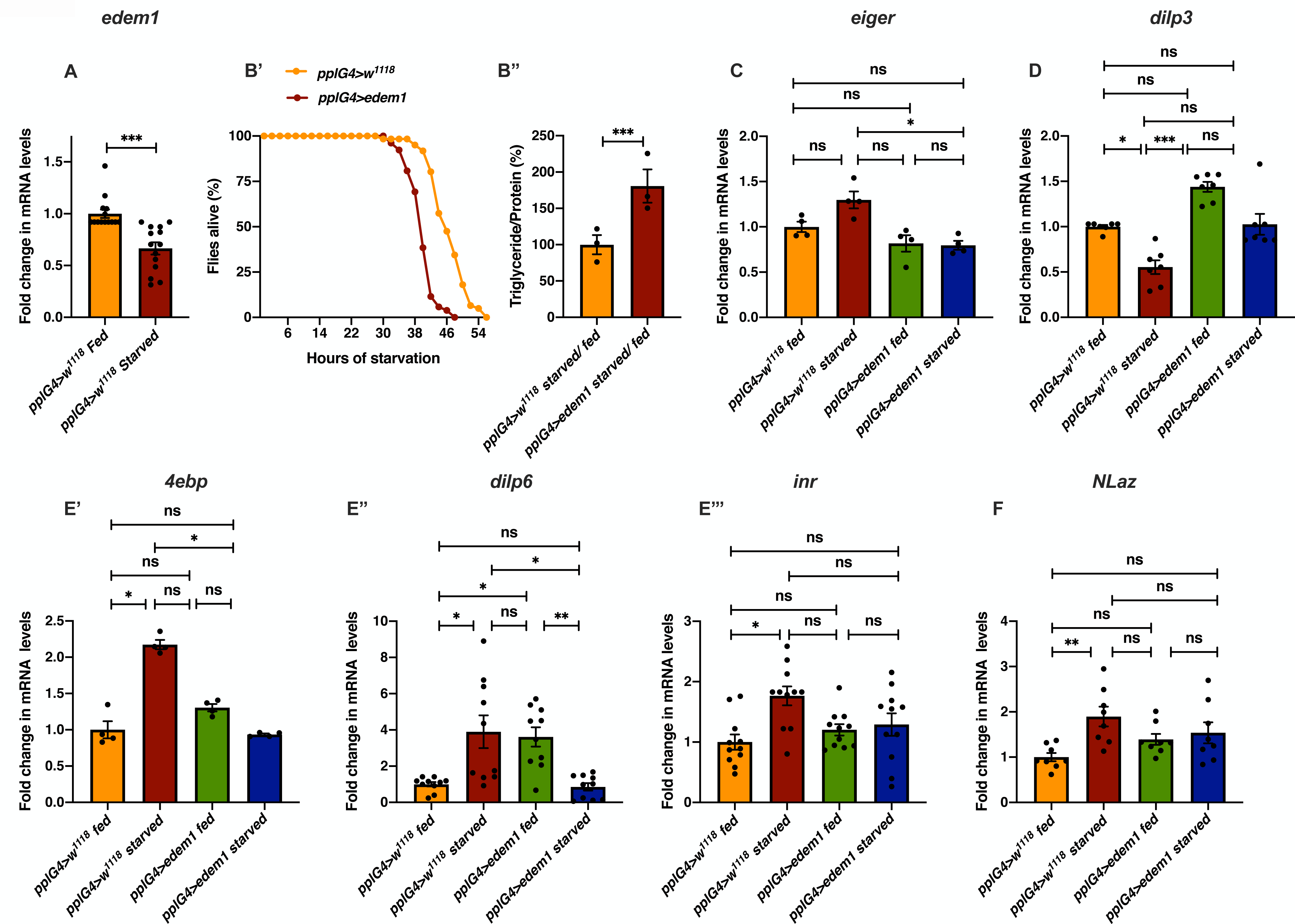
Reduction in *edem1* levels during starvation is crucial for survival. (A) Fold change in the mRNA levels of *edem1* upon starvation in *pplGal4>w^1118^* larvae. Values are normalised to *pplGal4>w^1118^* fed control and changes in the control starved are shown. [independent biological replicates = 14, P-value between control fed and starved larvae is <0.0001 (Mann-Whitney test)]. (B) Over-expression of *edem1* leads to enhanced sensitivity to starvation (Fig. 7B’), shown are percentage of male flies which were alive at various time points of starvation in the following genotypes - *pplGal4>w^1118^* and *pplGal4>UAS-edem1* [independent biological replicates = 3, number of flies used for control is 74, for *pplGal4>UAS-edem1* is 69. P-value between control and *UAS-edem1* is <0.0001 (Log-rank test), Wald test = 20.67 on df = 1, p < 0.001 (cox-proprtional hazard analysis)]. Fig. 7B’’ shows percentage reduction of triglyceride levels upon starvation. Data is shown as % ratio of starved to fed triglyceride to total protein levels, values are normalised to *pplGal4>w^1118^* and fold change in *pplGal4>UAS-edem1* is shown. [independent biological replicates = 3, P-value between control and *UAS-edem1* is 0.0001 (Unpaired t test).] (C) Overexpression of *edem1* in the fat body led to decreased levels of *eiger* mRNA when compared to control starved flies. Data is shown as fold change in mRNA levels, values are normalised to *pplGal4>w^1118^* and fold change in *pplGal4>w^1118^* starved and *pplGal4>UAS-edem1* fed and starved is shown. [independent biological replicates = 4. P-value between control fed and starved is >0.9999, P-value between control fed and *UAS-edem1* fed is >0.9999, P-value between control fed and *UAS-edem1* starved is 0.7133 and P-value between control starved and *UAS-edem1* fed is 0.0561, P-value between control starved and *UAS-edem1* starved is 0.0227, P-value between *UAS-edem1* fed and *UAS-edem1* starved is >0.9999 (Kruskal-Wallis test followed by Dunn’s post-hoc test)]. (D) Overexpression of *edem1* in the fat body increased the *dilp3* mRNA levels when compared to control starved flies. Data is shown as fold change in mRNA levels, values are normalised to *pplGal4>w^1118^* and fold change in *pplGal4>w^1118^* starved and *pplGal4>UAS-edem1* fed and starved is shown. [independent biological replicates = 7. P-value between control fed and starved is 0.0458, P-value between control fed and *UAS-edem1* fed is 0.4411, P-value between control fed and *UAS-edem1* starved is >0.9999 and P-value between control starved and *UAS-edem1* fed is <0.001, P-value between control starved and *UAS-edem1* starved is 0.2422, P-value between *UAS-edem1* fed and *UAS-edem1* starved is 0.0963 (Kruskal-Wallis test followed by Dunn’s post-hoc test)]. (E) Overexpression of *edem1* in the fat body led to decreased levels of *4ebp* (Fig. E’), *dilp6* (Fig. E’’) and *inr* (Fig. E’’’) when compared to control starved flies. Data is shown as fold change in mRNA levels, values are normalised to *pplGal4>w^1118^* and fold change in *pplGal4>w^1118^* starved and *pplGal4>UAS-edem1* fed and starved is shown. [independent biological replicates = 4 for *4ebp*, independent biological replicates = 10 for *dilp6* and independent biological replicates = 11 for *inr*. P-value between control fed and starved is 0.0178 for *4ebp*, 0.0218 for *dilp6* and 0.0051 for *inr*. P-value between control fed and *UAS-edem1* fed is 0.6193 for *4ebp*, 0.0102 for *dilp6* and >0.9999 for *inr*. P-value between control fed and *UAS-edem1* starved is >0.9999 for *4ebp*, >0.9999 for *dilp6* and 0.8366 for *inr*. P-value between control starved and *UAS-edem1* fed is >0.9999 for *4ebp*, >0.9999 for *dilp6* and 0.1931 for *inr*. P-value between control starved and *UAS-edem1* starved is 0.0286 for *4ebp*, 0.0132 for *dilp6* and 0.3774 for *inr*. P-value between *UAS-edem1* fed and *UAS-edem1* starved is 0.8249 for *4ebp*, 0.0060 for *dilp6* and >0.9999 for *inr* (Kruskal-Wallis test followed by Dunn’s post-hoc test)]. (F) Overexpression of *edem1* in the fat body led to decreased levels of *nlaz* mRNA when compared to control starved flies. Data is shown as fold change in mRNA levels, values are normalised to *pplGal4>w^1118^* and fold change in *pplGal4>w^1118^* starved and *pplGal4>UAS-edem1* fed and starved is shown. [independent biological replicates = 8. P-value between control fed and starved is 0.0052, P-value between control fed and *UAS-edem1* fed is 0.3727, P-value between control fed and *UAS-edem1* starved is 0.2410 and P-value between control starved and *UAS-edem1* fed is 0.8563, P-value between control starved and *UAS-edem1* starved is >0.9999, P-value between *UAS-edem1* fed and *UAS-edem1* starved is >0.9999 (Kruskal-Wallis test followed by Dunn’s post-hoc test)] **[*P-value *<0.05; ** <0.01,*** <0.001; Data information: In (A and B’’-F) data are presented as mean ± SEM*].**

## Discussion

Fluctuations in nutrient levels would trigger organism-wide changes, which includes alterations to various metabolic pathways. Changes in the metabolic pathways would aid the organism in managing the growth and maintenance of nutrient stores according to the availability of food. Apart from these biochemical changes, hunger elicits stereotypic behavioral responses, which includes an enhanced urge to feed, increased foraging, acceptance of unpalatable food, etc (Chouhan et al., 2017). Several of the crucial responses like mobilisation of stored nutrients and enhanced urge to feed, which aids the organism to survive nutrient deprivation is triggered by the reduction of systemic insulin signalling (Arsic & Guerin, 2008; Britton et al., 2002; Broughton et al., 2005; DiAngelo & Birnbaum, 2009; Kannan & Fridell, 2013; Rulifson et al., 2002; Tettweiler et al, 2005a). In *Drosophila,* IPCs respond to changes in the availability of food and modulate DILP levels under the control of the fat body, which acts as a nutrient sensor. Various fat body derived signals act on IPCs directly or indirectly, and the regulation of these signals in response to changes in the nutrient status of *Drosophila* plays a key role in maintaining systemic insulin levels (Agrawal et al., 2016; Bai et al., 2012; Colombani et al., 2003; Delanoue et al., 2016; Droujinine & Perrimon, 2016; Geminard et al., 2009; Ghosh & O’Connor, 2014; Koyama & Mirth, 2016; Rajan & Perrimon, 2012; Sano et al, 2015a; Sun et al., 2017). Here, we report a novel means by which the activity of a fat body derived signal on IPCs is regulated.

While investigating the mechanisms that function in the fat body to control *Drosophila* IPCs, we identified Edem1, an ER-resident protein involved in ERAD mediated protein quality control. Edem1 in the fat body maintains the activity of *Drosophila* TNFα Eiger and controls JNK signalling, thereby promoting normal IPC function, maintain systemic insulin signalling and metabolic homeostasis (Figs. 3 and 4). Eiger is activated by TACE, which cleaves the transmembrane form of Eiger and releases a soluble active form of Eiger into the hemolymph (Agrawal et al., 2016). TOR kinase, a key nutrient sensor, has been reported to control *tace* transcript levels and thereby Eiger activation. During low protein diets, due to reduced TOR signalling, fat body releases the soluble form of Eiger, which would act on IPCs and activate JNK signalling to regulate *dilp* gene expression. Here, we identify Edem1 as a regulator of Eiger through the control of *eiger* and *tace* gene expression (Fig. 3). We also show that activation of TOR signalling blocked the effects of suppression of *edem1* levels in the fat body (Fig. 5), substantiating the role of Edem1 in regulating Eiger activity. At the moment, it is not clear if the TOR pathway acts through Edem1 to regulate *eiger and tace* gene expression, thereby manage Eiger activity. More efforts are also needed to identify the exact molecular mechanism by which Edem1 regulates Eiger. We also found that the levels of Upd2, another FDS, is regulated by Edem1 function in the fat body. However, we observed that Upd2 does not act through IPCs and may act through another tissue unknown at the moment to regulate nutrient homeostasis.

Edem1 function in the fat body maintains systemic insulin signalling, and reduction of *edem1* levels in the fat body resulted in low systemic insulin signalling in larvae, which led to metabolic phenotypes as seen on circulating sugar levels and enhanced feeding in larvae; lipid and glycogen stores, enhanced resistance to starvation and increase in life span in adult flies (Fig 1). Low insulin signalling has been reported to cause these phenotypes by previous studies (Bai et al., 2012; Bohni et al., 1999; Broughton et al., 2005; Gronke et al., 2010a; Haselton et al., 2010a; Hong et al., 2012; Rulifson et al., 2002; Shingleton et al., 2005; Slaidina et al., 2009; Tatar et al., 2001; Teleman et al., 2006; Varghese et al., 2010a; Wu et al., 2005). We also show that the impact of reducing *edem1* levels on insulin signalling is due to the accumulation of DILP2 protein in the IPCs (Fig. 2C) and reduced *dilp3* transcript levels in the larval IPCs (Fig. 2B). However, it should be noted that we did not observe any changes at the protein and mRNA levels of other mNSC DILPs. DILPs are known to be regulated in a context specific manner, gene expression as well as protein levels in IPCs vary based on nutritional cues, developmental stages and various neural and endocrine signals that act on the IPCs (Gronke et al., 2010a; Hallier et al, 2016; Hong et al., 2012; Ikeya et al., 2002; Kim & Neufeld, 2015; Luo et al, 2014; Soderberg et al, 2012; Varghese et al, 2010b). Eiger activity on IPCs in response to low protein diet has been shown to suppress *dilp2* and *dilp5* transcript levels (Agrawal et al., 2016). Whereas, here we show that enhanced Eiger levels due to suppression of Edem1 expression in fed conditions affects *dilp3* transcription and DILP2 protein accumulation in the IPCs. Strictly the roles of individual DILPs are not understood, however, the effects of ablating IPCs, on growth and metabolism could be rescued by DILP2 expression alone (Haselton et al, 2010b; Rulifson et al., 2002). Many reports hint at effects on insulin signalling caused by an individual DILP or more than one DILP (Bai et al., 2012; Sudhakar et al, 2019).

Managing insulin signalling during nutrient withdrawal is crucial for mobilization of nutrient stores and survival. In response to starvation, we report that *edem1* transcripts are reduced (Fig. 7A). Reduction of *edem1* transcripts during starvation enhances *eiger* and *Nlaz* levels, which aids in lowering insulin signalling (Fig. 7C-F). This helps the flies to survive acute nutrient deprivation by mobilising energy stores (Fig. 7B and Fig. 8). Thus, Edem1 regulation plays an important role in eliciting responses to starvation and enhancing Edem1 levels affected survival during starvation, probably due to a failure in reducing insulin signalling and triglyceride mobilisation (Fig. 7B, 7D-E). Moreover, reducing Edem1 levels in fed conditions led to enhanced feeding responses, similar to starvation conditions, further suggesting an active role for Edem1 in survival against food deprivation (Fig. 1G). However, it is not yet clear if the function of Edem1 in regulating Eiger activity in the IPCs and systemic insulin signalling has any links to the ER stress pathway. Reduced *edem1* levels during starvation could be an outcome of reduced ER stress in response to low protein synthesis. Furthermore, reduction of *edem1* could cause aggregation of misfolded proteins in the ER, which might be responsible for the changes we report here on Eiger levels. However, blocking few other essential components of ERAD mechanism did not give us any expected results (Fig S3B). Hence, we are currently not sure whether Edem1 activity on managing systemic insulin signalling and nutrient homeostasis is linked to its ERAD functions.

**Fig. 8.**
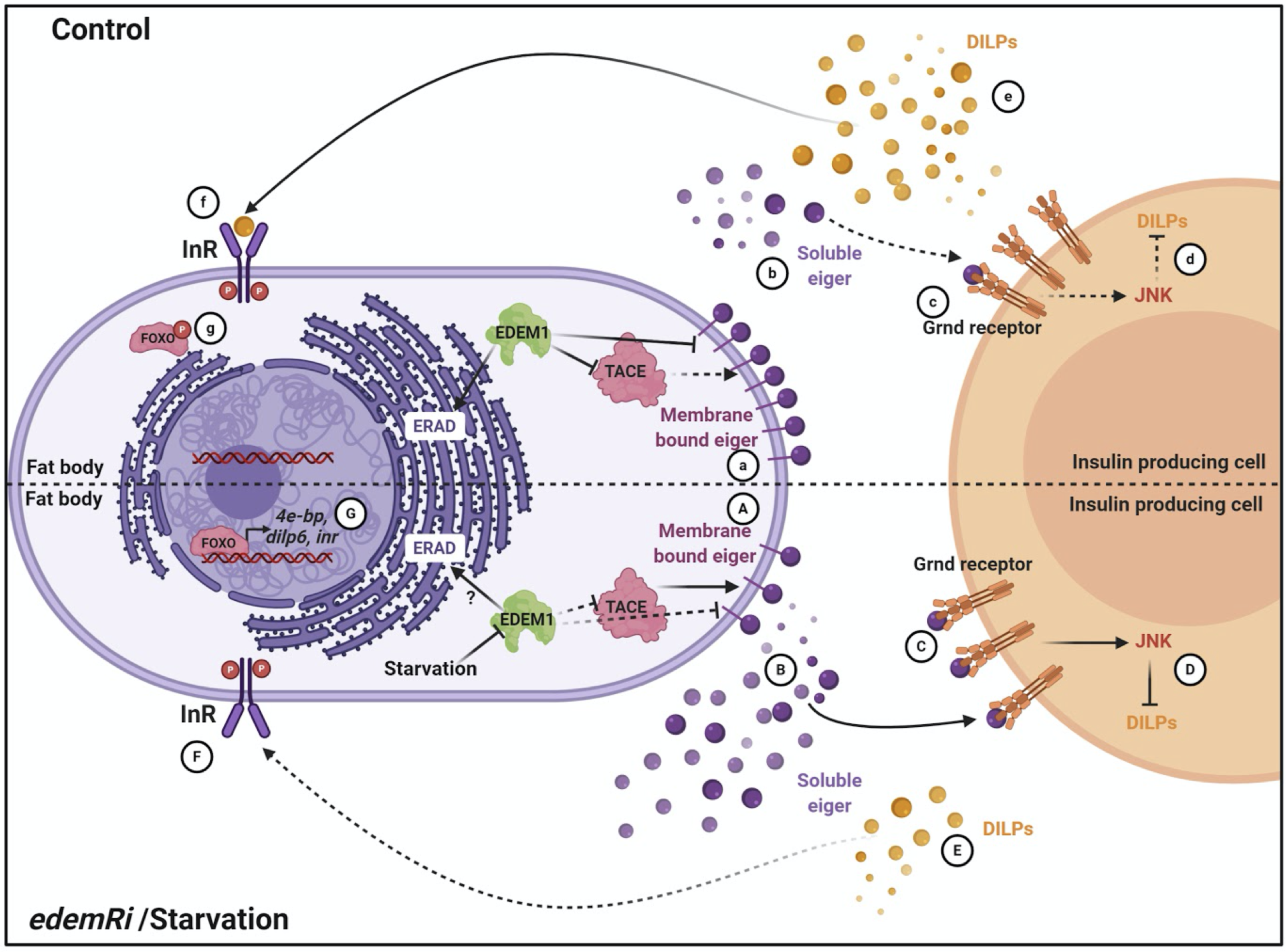
Working model. In control and fed conditions Edem1 blocks *tace* and *eiger* gene expression (a) and inhibits Eiger release (b). This maintains the insulin signalling (d-g) as JNK signalling is kept low in the IPCs (c). In starved conditions and in the *edem1*Ri background, *edem1* levels are low, and *tace* and *eiger* gene expression is increased (A) and leading to increased Eiger secretion into the hemolymph (B). Soluble Eiger binds to the Grnd receptors in the IPCs and activate JNK signalling which inhibits insulin signalling (C and D). Reduction of insulin signalling mediated by reduction of Edem1 in the fat body (E-G) aid in survival of flies. (Created with BioRender.com)

To summarize, we show that Edem1, a key ERAD regulator, aids in the maintenance of nutrient homeostasis by managing the activity of TNFα Eiger on Drosophila insulin producing cells (Fig 8). During fed conditions Edem1 suppress Eiger levels, which allows optimal insulin signalling and maintain a steady metabolic status. In response to starvation, suppression of Edem1 leads to a reduction of insulin signalling and mobilisation of energy reserves, which aids in survival during acute food deprivation.

## Materials and Methods

### Fly strains

Fly stocks were reared in vials with standard food which consisted of 5.8% cornmeal, 5% dextrose, 2.36% yeast, 0.8% agar and 10% Nipagen in 100% ethanol. All the flies were maintained at 25 °C with 12 h:12 h light:dark cycle. *UAS-InR^A1325D^* (RRID:BDSC_8263) and *UAS-edem1-RNAi* (RRID:BDSC_58298) were obtained from Bloomington Drosophila stock center (BDSC). The RNAi lines used were obtained from Vienna Drosophila resource center (VDRC): *UAS-edem1-RNAi* (stock# 6923, 6922), *UAS-eiger-RNAi* (stock# 45253), *UAS-grnd-RNAi* (stock # 43454), *UAS-dilp6-RNAi* (GD) (stock# 45218), *UAS-herp-RNAi* (stock# 11724, 11725) and *UAS-sip3-RNAi* (stock# 6870, 107060). *dilp2*-Gal4/ *CyO*GFP, *pumpless*-Gal4 and *w^1118^* were obtained from Stephen Cohen. *UAS-dEDEM1* was from Koichi Iijima, *UAS-rheb* was obtained from Jagat. K. Roy, *UAS-upd2-EGFP/ TM3Sb* and 10XSTAT92E-GFP were obtained from Akhila Rajan. In order to match the genetic background all the fly strains used in this study were back-crossed into an isogenic *w^1118^* background for at least 6 generations.

### Triglyceride and glycogen measurements

All experiments were carried out in controlled growth conditions as described here, unless mentioned otherwise. Fifty 1^st^ in-star larvae were collected in fresh food vials avoiding overcrowding within 2–3 h of hatching. GFP balancers were used wherever required to aid in genotyping. Freshly emerged adult male flies were collected (15 per vial) and 5-day old flies were used for triglyceride and glycogen measurement unless mentioned otherwise. 5 flies in triplicates per genotype were homogenized in 0.05% Tween-20 using Bullet Blender Storm BBY24M from Next Advance. Each experiment was replicated independently, number of independent biological replicates is mentioned for each experiment in the figure legends. The homogenate was heat inactivated at 70 °C for 5 min and then centrifuged at 14000 rpm for 3 min. Serum triglyceride determination kit (Cat. # TR0100) from Sigma was used to quantify triglyceride levels and protein levels were measured using the Quick Start™ Bradford 1X Dye Reagent (Cat. # 500-0205) from Biorad. This was followed by colorimetric estimation using TECAN Infinite M200 pro-multimode plate reader in 96-well format. The absorption maximum of 540 nm and 595 nm were used for triglyceride and protein content respectively. Sample preparation for glycogen measurement was similar to triglycerides, following the manufacturer’s protocol (Cat. # MAK016 from Sigma). The absorbance was measured at 570 nm. For triglyceride and glycogen utilization assay, 5 day old males (15 per vial) were transferred to vials containing 1% agar, were collected at the indicated time points and homogenized as mentioned above. Each experiment was replicated independently, number of replicates (n) is mentioned for each experiment in the figure legends.

### Starvation sensitivity assay

Fifty 1^st^ in-star larvae were collected in fresh food vials avoiding overcrowding within 2–3 h of hatching. GFP balancers were used wherever required to aid in genotyping. Freshly emerged adult male flies were collected (15 per vial). For starvation sensitivity assay, 15 (5-day old) male flies were transferred to vials containing 1% agar and the number of dead flies was counted every 2 hours. Multiple vials were set as technical replicates. These experiments were replicated independently and number of independent biological replicates are mentioned in the figure legends.

### Lifespan assay

Adult lifespan assay was estimated with data obtained from three independent biological replicates for each genotype. Fifty 1^st^ in-star larvae were collected in fresh food vials and freshly emerged adult male flies were collected (15 per vial). Multiple vials of adult male flies were set as technical replicates. These flies were flipped into fresh media every 2 days and the dead flies and the escapers were scored.

### Glucose assay

Fifty 1^st^ in-star larvae were collected in fresh food vials. Larvae at 3^rd^ in-star stage (five larvae for every prep) larvae were used to isolate hemolymph using Zymo-Spin™ IIIC (C1006-250) from Zymo Research. 1 μl of hemolymph was diluted to 50 μl with autoclaved milli-Q water. 100 μl of glucose assay reagent (Cat. # GAGO20) from Sigma was added and the reaction was incubated at 37 °C for 30 min. The reaction was stopped with 100 μl of 12 NH_2_SO_4_. The glucose content was analyzed using colorimetric quantification at 540 nm using TECAN Infinite M200 pro-multi-mode plate reader in 96-well format. Hemolymph glucose measurements were replicated independently and number of replicates are mentioned in the figure legends.

### Larval starvation

Fifty 1^st^ in-star larvae were collected in fresh food vials. Third in-star non-crawling larvae of the desired genotypes were kept for starvation on 1% agar vials for 12hrs, after washing them with milli-Q water to make sure that there were no traces of media left behind. After 12 hrs the larvae were plunged for the qPCR experiments. The starvation experiments were replicated independently and number of replicates are mentioned in the figure legends.

### Feeding assay

Fifty 1^st^ in-star larvae were collected in fresh food vials. Larvae at 3^rd^ in-star stage (10 each) or 5-day old flies (5 each) were fed for 3 h or 30 min respectively, with colored food with Orange G dye (Cat. # 1936-15-8) from Sigma. The larvae/flies were homogenized using 0.05% Tween-20. The homogenate was analyzed colorimetrically at 492 nm using TECAN Infinite M200 pro-multi-mode plate reader in 96-well format. The absorbance of the homogenate was directly proportional to the food intake. The feeding experiments were replicated independently and number of replicates are mentioned in the figure legends.

### Quantitative RT-PCR

Fifty 1^st^ in-star larvae were collected in fresh food vials. Third in-star wandering larvae for each genotype were collected and were flash-frozen. For fly qRT-PCR experiments with, the flies were starved for 24 h before flash freezing them and two sets of RNAs were isolated independently from the heads and the body. These experiments were replicated independently and number of replicates (n) are mentioned in the figure legends. Total RNA was isolated with QIAGEN RNeasy Plus Mini Kit (Cat. # 74134) and was quantified using Qubit™ RNA HS Assay Kit (Cat. # Q32852). An equal amount of RNA from each sample was reverse transcribed using SuperScript® III First-Strand Synthesis System (Cat. # 18080051) from ThermoFisher Scientific. Quantitative RT-PCR was performed using Bio-Rad CFX96™ with the cDNA template, Power SYBR® Green PCR Master Mix (Cat. # 4368702) from ThermoFisher Scientific and a primer concentration of 312.5 nM. The data were normalized to *rp49*. The sequences of the primers used are mentioned in Table 1.

**Table 1.**
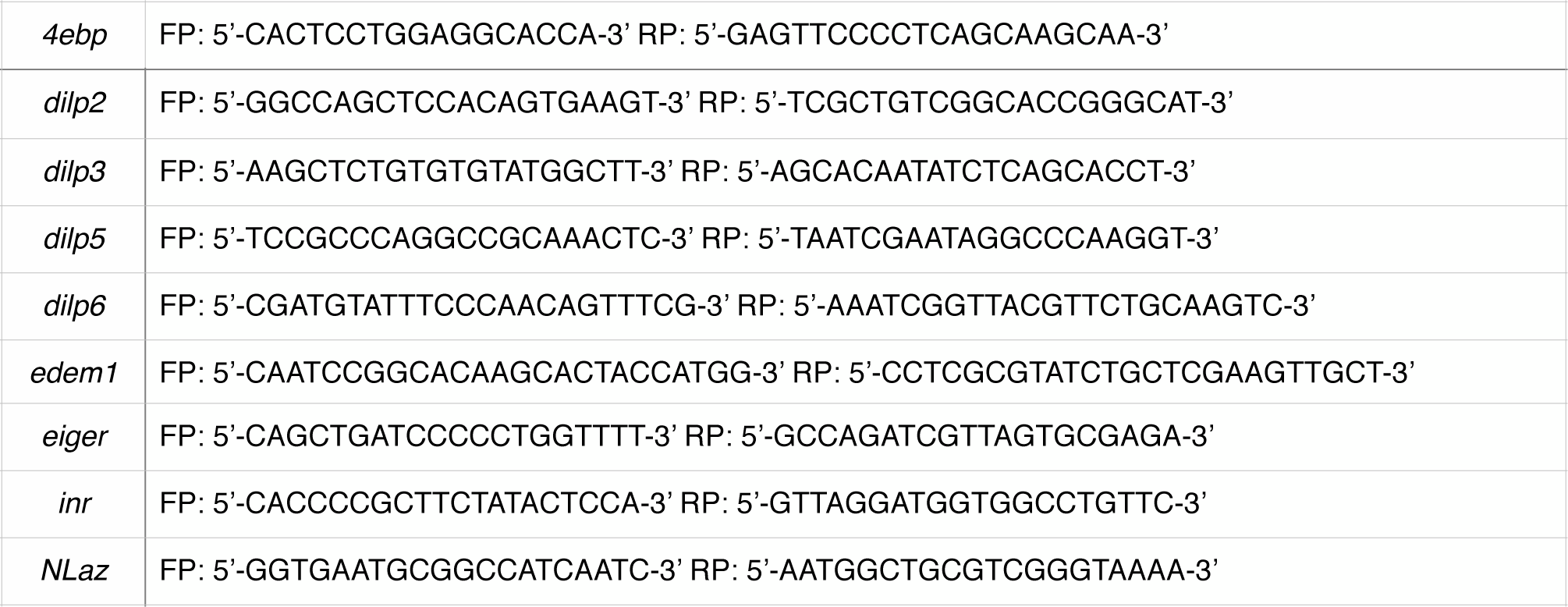
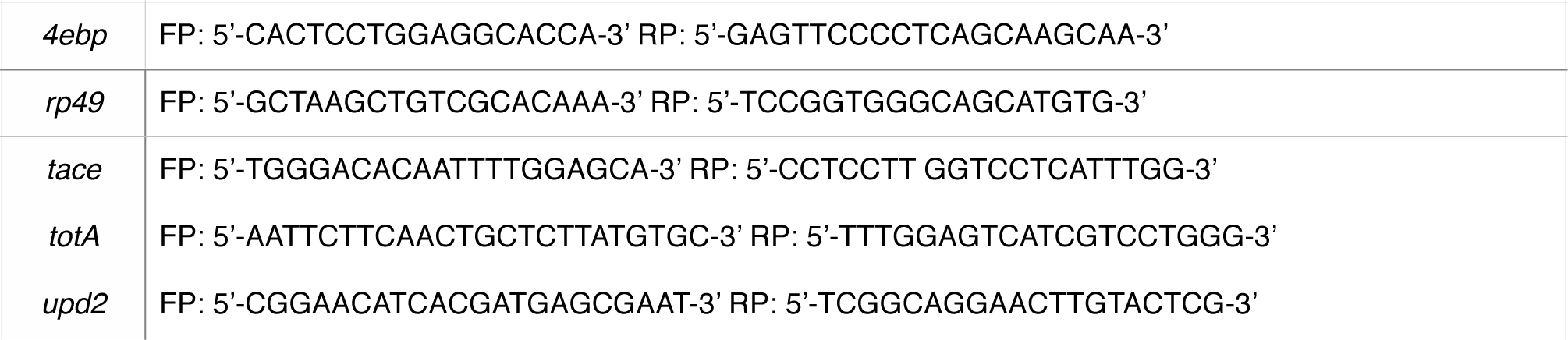
List of primer sequences used in the manuscript.

### Immunohistochemistry

DILP2 peptide corresponding to the sequence TRQRQGIVERC (amino acids 108-118) was used as an immunogen to raise DILP2 polyclonal antibody in rabbit (Eurogentec, Belgium). Mouse anti-GFP (Cat. # 632375 from Living Colors) was used. About 10 larvae (3^rd^ in-star wandering) were used to dissect the brains in ice-cold 1X phosphate buffered saline PBS (Cat. # P4417 from Sigma) per genotype for each experiment. The dissections were repeated independently and number of replicates (n) are mentioned in the figure legends. The tissue samples were fixed using 4% paraformaldehyde (PFA) (Cat # P6148 from Sigma) at room temperature for 20 min. PFA was removed and the tissues were washed with 1X phosphate buffered saline + 0.1% Triton X-100 (Cat. # 161-0407 from Bio-Rad) PBT. Blocking solution [PBT+ 0.1% bovine serum albumin BSA (Cat. # A2153 from Sigma) BBT] was added to the tissues and the tissues were incubated at room temperature for 45 min. Primary antibody against DILP2 and GFP were diluted in BBT in 1:1000 and 1:500 dilutions respectively. The samples were incubated with primary antibody overnight at 4 °C with constant rotation. Then the tissues were washed extensively with 1X PBT and incubated with secondary antibody at room temperature for 2 h. The secondary antibodies, Alexa Fluor® 488 Goat Anti-Rabbit IgG (Cat. # A27034) and Alexa Fluor® 633 Goat Anti-Mouse IgG (Cat. # A-21050) were diluted in 1:500 dilution in BBT. After 2 h the samples were washed extensively and mounted with a drop of SlowFade® Gold Antifade Reagent with DAPI (Cat. # S36939) from ThermoFisher Scientific. The tissues were imaged using a Leica DM6000B upright microscope and processed using ImageJ software. Corrected total cell fluorescence (CTCF) was calculated using the formula CTCF = Integrated Density – (Area of selected cell X Mean fluorescence of background readings).

### *ex-vivo* organ co-culture

For the *ex-vivo* organ co-culture, larval hemolymph was isolated from 10 (3^rd^ in-star crawling) larvae and was incubated with 10 brains from 3^rd^ in-star crawling larvae of the desired genotypes in Shields and Sang medium (Cat. # S3652 from Sigma) at room temperature for 2 h with constant shaking. The larval brains were then fixed in 4 % PFA and stained for DILP2 as mentioned above and imaged. For the qRT-PCR, larval hemolymph was isolated from 35 (3^rd^ in-star crawling) larvae and was incubated with 35 brains from 3^rd^ in-star crawling larvae of the desired genotype. After 2 h, the brains were flash frozen and the RNA was isolated as mentioned before.

### Western blotting

5-day old adult flies 5 each of the desired genotype were homogenised in 40 μl RIPA buffer with cOmplete^™^, EDTA-free Protease Inhibitor Cocktail (Cat # 4693132001 from Sigma). The homogenates were centrifuged at full speed. The samples were denatured in 2x Laemmli sample buffer (Cat # 1610737 from Biorad) at 95°C and run on 10% SDS-PAGE. Immobilon Western Chemiluminescent HRP Substrate (Cat # WBKLS0050 from Merck, Millipore) was used for antibody detection after blotting on PVDF membrane (Cat # 162-0177 from Biorad). The following primary antibodies were used: anti-Egr 1:50 (generous gift from Konrad Basler) and anti-actin 1:3000 (Cat # 612656 from BD Biosciences). For quantification, the intensity of soluble eiger protein bands was normalized to the intensity of actin bands using Image J software.

### Statistical analysis

All the experiments were done in biological replicates as indicated and the error bars represent the standard error mean (SEM). The graphs were plotted using GraphPad Prism8 software. Significance was tested using Unpaired t-test, Welch test, Mann-Whitney test and Kruskal-Wallis test (followed by Dunn’s post hoc test) with * representing p-value < 0.05, ** p-value < 0.01, *** p-value < 0.001. The starvation and lifespan datasets were subjected to Log rank test followed by cox-proportional hazard analysis using R software to analyze the trends in the survival of flies in both the assays.

## Acknowledgements

We thank Nisha N. Kannan and Smitha Vishnu for critical comments on this manuscript. HP was supported by IISER TVM PHD fellowship. JV was supported by Ramanujan fellowship, DST, India. This work was also supported by generous funds from SERB, DST, India and core funding from IISER TVM, MHRD, India.

## Authors contributions

HP and JV conceptualized the paper. HP performed research and analysed data. HP and JV wrote the paper. JV acquired funding and supervised research.

## Conflict of interest

The authors declare no conflict of interest

## Notes

### Competing Interest Statement

The authors have declared no competing interest.

### Summary of Updates

New data added, few changes to the text and figures

## References

1. Accili D, Drago J, Lee EJ, Johnson MD, Cool MH, Salvatore P, Asico LD, Jose PA, Taylor SI, Westphal H (1996) Early neonatal death in mice homozygous for a null allele of the insulin receptor gene. Nat Genet 12: 106–109

2. Agrawal N, Delanoue R, Mauri A, Basco D, Pasco M, Thorens B, Leopold P (2016) The Drosophila TNF Eiger Is an Adipokine that Acts on Insulin-Producing Cells to Mediate Nutrient Response. Cell Metabolism 23: 675–684

3. Andersen DS, Colombani J, Palmerini V, Chakrabandhu K, Boone E, Rothlisberger M, Toggweiler J, Basler K, Mapelli M, Hueber AO et al (2015) The Drosophila TNF receptor Grindelwald couples loss of cell polarity and neoplastic growth. Nature 522: 482-+

4. Araki K, Nagata K (2011) Protein folding and quality control in the ER. Cold Spring Harb Perspect Biol 3: a007526

5. Arsic D, Guerin PM (2008) Nutrient content of diet affects the signaling activity of the insulin/target of rapamycin/p70 S6 kinase pathway in the African malaria mosquito Anopheles gambiae. J Insect Physiol 54: 1226–1235

6. Bai H, Kang P, Hernandez AM, Tatar M (2013) Activin signaling targeted by insulin/dFOXO regulates aging and muscle proteostasis in Drosophila. Plos Genet 9: e1003941

7. Bai H, Kang P, Tatar M (2012) Drosophila insulin-like peptide-6 (dilp6) expression from fat body extends lifespan and represses secretion of Drosophila insulin-like peptide-2 from the brain. Aging Cell 11: 978–985

8. Bai H, Post S, Kang P, Tatar M (2015) Drosophila Longevity Assurance Conferred by Reduced Insulin Receptor Substrate Chico Partially Requires d4eBP. PLoS One 10: e0134415

9. Bohni R, Riesgo-Escovar J, Oldham S, Brogiolo W, Stocker H, Andruss BF, Beckingham K, Hafen E (1999) Autonomous control of cell and organ size by CHICO, a Drosophila homolog of vertebrate IRS1-4. Cell 97: 865–875

10. Bonafe M, Barbieri M, Marchegiani F, Olivieri F, Ragno E, Giampieri C, Mugianesi E, Centurelli M, Franceschi C, Paolisso G (2003) Polymorphic variants of insulin-like growth factor I (IGF-I) receptor and phosphoinositide 3-kinase genes affect IGF-I plasma levels and human longevity: Cues for an evolutionarily conserved mechanism of life span control. J Clin Endocr Metab 88: 3299–3304

11. Britton JS, Lockwood WK, Li L, Cohen SM, Edgar BA (2002) Drosophila’s insulin/PI3-kinase pathway coordinates cellular metabolism with nutritional conditions. Dev Cell 2: 239–249

12. Brogiolo W, Stocker H, Ikeya T, Rintelen F, Fernandez R, Hafen E (2001) An evolutionarily conserved function of the Drosophila insulin receptor and insulin-like peptides in growth control. Curr Biol 11: 213–221

13. Broughton SJ, Piper MD, Ikeya T, Bass TM, Jacobson J, Driege Y, Martinez P, Hafen E, Withers DJ, Leevers SJ et al (2005) Longer lifespan, altered metabolism, and stress resistance in Drosophila from ablation of cells making insulin-like ligands. Proc Natl Acad Sci U S A 102: 3105–3110

14. Broughton SJ, Slack C, Alic N, Metaxakis A, Bass TM, Driege Y, Partridge L (2010) DILP-producing median neurosecretory cells in the Drosophila brain mediate the response of lifespan to nutrition. Aging Cell 9: 336–346

15. Chouhan NS, Wolf R, Heisenberg M (2017) Starvation promotes odor/feeding-time associations in flies. Learn Memory 24: 318–321

16. Clancy DJ, Gems D, Harshman LG, Oldham S, Stocker H, Hafen E, Leevers SJ, Partridge L (2001) Extension of life-span by loss of CHICO, a Drosophila insulin receptor substrate protein. Science 292: 104–106

17. Colombani J, Raisin S, Pantalacci S, Radimerski T, Montagne J, Leopold P (2003) A nutrient sensor mechanism controls Drosophila growth. Cell 114: 739–749

18. Delanoue R, Meschi E, Agrawal N, Mauri A, Tsatskis Y, McNeill H, Leopold P (2016) Drosophila insulin release is triggered by adipose Stunted ligand to brain Methuselah receptor. Science 353: 1553–1556

19. DiAngelo JR, Birnbaum MJ (2009) Regulation of Fat Cell Mass by Insulin in Drosophila melanogaster. Mol Cell Biol 29: 6341–6352

20. Droujinine IA, Perrimon N (2016) Interorgan Communication Pathways in Physiology: Focus on Drosophila. Annu Rev Genet 50: 539–570

21. Erion R, Sehgal A (2013) Regulation of insect behavior via the insulin-signaling pathway. Front Physiol 4

22. Fabrizio P, Pozza F, Pletcher SD, Gendron CM, Longo VD (2001) Regulation of longevity and stress resistance by Sch9 in yeast. Science 292: 288–290

23. Fernandez AM, Torres-Aleman I (2012) The many faces of insulin-like peptide signalling in the brain. Nat Rev Neurosci 13: 225–239

24. Garami A, Zwartkruis FJT, Nobukuni T, Joaquin M, Roccio M, Stocker H, Kozma SC, Hafen E, Bos JL, Thomas G (2003) Insulin activation of Rheb, a mediator of mTOR/ S6K/4E-BP signaling, is inhibited by TSC1 and 2. Mol Cell 11: 1457–1466

25. Geminard C, Rulifson EJ, Leopold P (2009) Remote control of insulin secretion by fat cells in Drosophila. Cell Metab 10: 199–207

26. Ghosh AC, O’Connor MB (2014) Systemic Activin signaling independently regulates sugar homeostasis, cellular metabolism, and pH balance in Drosophila melanogaster. P Natl Acad Sci USA 111: 5729–5734

27. Giannakou ME, Goss M, Junger MA, Hafen E, Leevers SJ, Partridge L (2004) Long-lived Drosophila with overexpressed dFOXO in adult fat body. Science 305: 361

28. Giannakou ME, Partridge L (2007) Role of insulin-like signalling in Drosophila lifespan. Trends Biochem Sci 32: 180–188

29. Gronke S, Clarke DF, Broughton S, Andrews TD, Partridge L (2010a) Molecular evolution and functional characterization of Drosophila insulin-like peptides. Plos Genet 6: e1000857

30. Gronke S, Clarke DF, Broughton S, Andrews TD, Partridge L (2010b) Molecular Evolution and Functional Characterization of Drosophila Insulin-Like Peptides. Plos Genet 6

31. Hallier B, Schiemann R, Cordes E, Vitos-Faleato J, Walter S, Heinisch JJ, Malmendal A, Paululat A, Meyer H (2016) Drosophila neprilysins control insulin signaling and food intake via cleavage of regulatory peptides. Elife 5

32. Haselton A, Sharmin E, Schrader J, Sah M, Poon P, Fridell YW (2010a) Partial ablation of adult Drosophila insulin-producing neurons modulates glucose homeostasis and extends life span without insulin resistance. Cell Cycle 9: 3063–3071

33. Haselton A, Sharmin E, Schrader J, Sah M, Poon P, Fridell YWC (2010b) Partial ablation of adult Drosophila insulin-producing neurons modulates glucose homeostasis and extends life span without insulin resistance. Cell Cycle 9: 3063–3071

34. Hirosumi J, Tuncman G, Chang LF, Gorgun CZ, Uysal KT, Maeda K, Karin M, Hotamisligil GS (2002) A central role for JNK in obesity and insulin resistance. Nature 420: 333–336

35. Holzenberger M, Dupont J, Ducos B, Leneuve P, Geloen A, Even PC, Cervera P, Le Bouc Y (2003) IGF-1 receptor regulates lifespan and resistance to oxidative stress in mice. Nature 421: 182–187

36. Hong SH, Lee KS, Kwak SJ, Kim AK, Bai H, Jung MS, Kwon OY, Song WJ, Tatar M, Yu K (2012) Minibrain/Dyrk1a Regulates Food Intake through the Sir2-FOXO-sNPF/NPY Pathway in Drosophila and Mammals. Plos Genet 8

37. Hull-Thompson J, Muffat J, Sanchez D, Walker DW, Benzer S, Ganfornina MD, Jasper H (2009) Control of Metabolic Homeostasis by Stress Signaling Is Mediated by the Lipocalin NLaz. Plos Genet 5

38. Hwangbo DS, Gershman B, Tu MP, Palmer M, Tatar M (2005) Drosophila dFOXO controls lifespan and regulates insulin signalling in brain and fat body (vol 429, pg 562, 2004). Nature 434: 118–118

39. Ikeya T, Galic M, Belawat P, Nairz K, Hafen E (2002) Nutrient-dependent expression of insulin-like peptides from neuroendocrine cells in the CNS contributes to growth regulation in Drosophila. Curr Biol 12: 1293–1300

40. Kahn SE, Hull RL, Utzschneider KM (2006) Mechanisms linking obesity to insulin resistance and type 2 diabetes. Nature 444: 840–846

41. Kannan K, Fridell YWC (2013) Functional implications of Drosophila insulin-like peptides in metabolism aging, and dietary restriction. Front Physiol 4

42. Katic M, Kahn CR (2005) The role of insulin and IGF-1 signaling in longevity. Cellular and Molecular Life Sciences 62: 320–343

43. Kenyon C, Chang J, Gensch E, Rudner A, Tabtiang R (1993) A C-Elegans Mutant That Lives Twice as Long as Wild-Type. Nature 366: 461–464

44. Kim J, Neufeld TP (2015) Dietary sugar promotes systemic TOR activation in Drosophila through AKH-dependent selective secretion of Dilp3. Nat Commun 6

45. Kimura KD, Tissenbaum HA, Liu YX, Ruvkun G (1997) daf-2, an insulin receptor-like gene that regulates longevity and diapause in Caenorhabditis elegans. Science 277: 942–946

46. Koyama T, Mirth CK (2016) Growth-Blocking Peptides As Nutrition-Sensitive Signals for Insulin Secretion and Body Size Regulation. Plos Biol 14

47. Kroeger H, Chiang WC, Lin JH (2012) Endoplasmic Reticulum-Associated Degradation (ERAD) of Misfolded Glycoproteins and Mutant P23H Rhodopsin in Photoreceptor Cells. Adv Exp Med Biol 723: 559–565

48. Liu CY, Kaufman RJ (2003) The unfolded protein response. J Cell Sci 116: 1861–1862

49. Liu JP, Baker J, Perkins AS, Robertson EJ, Efstratiadis A (1993) Mice carrying null mutations of the genes encoding insulin-like growth factor I (Igf-1) and type 1 IGF receptor (Igf1r). Cell 75: 59–72

50. Luo J, Lushchak OV, Goergen P, Williams MJ, Nassel DR (2014) Drosophila insulin-producing cells are differentially modulated by serotonin and octopamine receptors and affect social behavior. PLoS One 9: e99732

51. Min KJ, Yamamoto R, Buch S, Pankratz M, Tatar M (2008) Drosophila lifespan control by dietary restriction independent of insulin-like signaling. Aging Cell 7: 199–206

52. Molinari M, Calanca V, Galli C, Lucca P, Paganetti P (2003) Role of EDEM in the release of misfolded glycoproteins from the calnexin cycle. Science 299: 1397–1400

53. Nassel DR (2012) Insulin-producing cells and their regulation in physiology and behavior of Drosophila. Can J Zool 90: 476–488

54. Oh SW, Mukhopadhyay A, Svrzikapa N, Jiang F, Davis RJ, Tissenbaum HA (2005) JNK regulates lifespan in Caenorhabditis elegans by modulating nuclear translocation of forkhead transcription factor/DAF-16. P Natl Acad Sci USA 102: 4494–4499

55. Oldham S (2011) Obesity and nutrient sensing TOR pathway in flies and vertebrates: Functional conservation of genetic mechanisms. Trends Endocrin Met 22: 45–52

56. Oldham S, Montagne J, Radimerski T, Thomas G, Hafen E (2000) Genetic and biochemical characterization of dTOR, the Drosophila homolog of the target of rapamycin. Gene Dev 14: 2689–2694

57. Partridge L, Alic N, Bjedov I, Piper MD (2011) Ageing in Drosophila: the role of the insulin/Igf and TOR signalling network. Exp Gerontol 46: 376–381

58. Pasco MY, Leopold P (2012) High Sugar-Induced Insulin Resistance in Drosophila Relies on the Lipocalin Neural Lazarillo. Plos One 7

59. Post S, Karashchuk G, Wade JD, Sajid W, De Meyts P, Tatar M (2018) Drosophila Insulin-Like Peptides DILP2 and DILP5 Differentially Stimulate Cell Signaling and Glycogen Phosphorylase to Regulate Longevity. Front Endocrinol (Lausanne) 9: 245

60. Puig O, Marr MT, Ruhf ML, Tjian R (2003) Control of cell number by Drosophila FOXO: downstream and feedback regulation of the insulin receptor pathway. Genes Dev 17: 2006–2020

61. Rajan A, Perrimon N (2012) Drosophila Cytokine Unpaired 2 Regulates Physiological Homeostasis by Remotely Controlling Insulin Secretion. Cell 151: 123–137

62. Rajan A, Perrimon N (2013) Drosophila Cytokine Unpaired 2 Regulates Physiological Homeostasis by Remotely Controlling Insulin Secretion (vol 151, pg 123, 2012). Cell 152: 1197–1197

63. Rulifson EJ, Kim SK, Nusse R (2002) Ablation of insulin-producing neurons in flies: Growth and diabetic phenotypes. Science 296: 1118–1120

64. Sano H, Nakamura A, Texada MJ, Truman JW, Ishimoto H, Kamikouchi A, Nibu Y, Kume K, Ida T, Kojima M (2015a) Correction: The Nutrient-Responsive Hormone CCHamide-2 Controls Growth by Regulating Insulin-like Peptides in the Brain of Drosophila melanogaster. Plos Genet 11: e1005481

65. Sano H, Nakamura A, Texada MJ, Truman JW, Ishimoto H, Kamikouchi A, Nibu Y, Kume K, Ida T, Kojima M (2015b) The Nutrient-Responsive Hormone CCHamide-2 Controls Growth by Regulating Insulin-like Peptides in the Brain of Drosophila melanogaster. Plos Genet 11

66. Saucedo LJ (2003) Rheb promotes cell growth as a compound of the insulin/TOR signalling network (vol 5, pg 566, 2003). Nat Cell Biol 5: 680–680

67. Shimokawa I, Higami Y, Tsuchiya T, Otani H, Komatsu T, Chiba T, Yamaza H (2003) Life span extension by reduction of the growth hormone-insulin-like growth factor-1 axis: relation to caloric restriction. FASEB J 17: 1108–1109

68. Shingleton AW, Das J, Vinicius L, Stern DL (2005) The temporal requirements for insulin signaling during development in Drosophila. Plos Biol 3: e289

69. Slaidina M, Delanoue R, Gronke S, Partridge L, Leopold P (2009) A Drosophila insulin-like peptide promotes growth during nonfeeding states. Dev Cell 17: 874–884

70. Soderberg JA, Carlsson MA, Nassel DR (2012) Insulin-Producing Cells in the Drosophila Brain also Express Satiety-Inducing Cholecystokinin-Like Peptide, Drosulfakinin. Front Endocrinol (Lausanne) 3: 109

71. Sonntag WE, Carter CS, Ikeno Y, Ekenstedt K, Carlson CS, Loeser RF, Chakrabarty S, Lee S, Bennett C, Ingram R et al (2005) Adult-onset growth hormone and insulin-like growth factor I deficiency reduces neoplastic disease, modifies age-related pathology, and increases life span. Endocrinology 146: 2920–2932

72. Sudhakar SR, Pathak H, Rehman N, Fernandes J, Vishnu S, Varghese J (2019) Insulin signalling elicits hunger-induced feeding in Drosophila. Dev Biol

73. Sun JH, Liu C, Bai XB, Li XT, Li JY, Zhang ZP, Zhang YP, Guo J, Li Y (2017) Drosophila FIT is a protein-specific satiety hormone essential for feeding control. Nat Commun 8

74. Tatar M, Kopelman A, Epstein D, Tu MP, Yin CM, Garofalo RS (2001) A mutant Drosophila insulin receptor homolog that extends life-span and impairs neuroendocrine function. Science 292: 107–110

75. Teleman AA, Maitra S, Cohen SM (2006) Drosophila lacking microRNA miR-278 are defective in energy homeostasis. Genes Dev 20: 417–422

76. Tettweiler G, Miron M, Jenkins M, Sonenberg N, Lasko PF (2005) Starvation and oxidative stress resistance in Drosophila are mediated through the eIF4E-binding protein, d4E-BP. Gene Dev 19: 1840–1843

77. Varghese J, Lim SF, Cohen SM (2010a) Drosophila miR-14 regulates insulin production and metabolism through its target, sugarbabe. Gene Dev 24: 2748–2753

78. Varghese J, Lim SF, Cohen SM (2010b) Drosophila miR-14 regulates insulin production and metabolism through its target, sugarbabe. Gene Dev 24: 2748–2753

79. Wang MC, Bohmann D, Jasper H (2005) JNK extends life span and limits growth by antagonizing cellular and organism-wide responses to insulin signaling. Cell 121: 115–125

80. Wu Q, Zhang Y, Xu J, Shen P (2005) Regulation of hunger-driven behaviors by neural ribosomal S6 kinase in Drosophila. Proc Natl Acad Sci U S A 102: 13289–13294

81. Zhang T, Branch A, Shen P (2013) Octopamine-mediated circuit mechanism underlying controlled appetite for palatable food in Drosophila. Proc Natl Acad Sci U S A 110: 15431–15436

