## Supplementary figures and legends for "Edem1 activity in the fat body regulates insulin signalling and metabolic homeostasis in *Drosophila*"

### Fig. S1

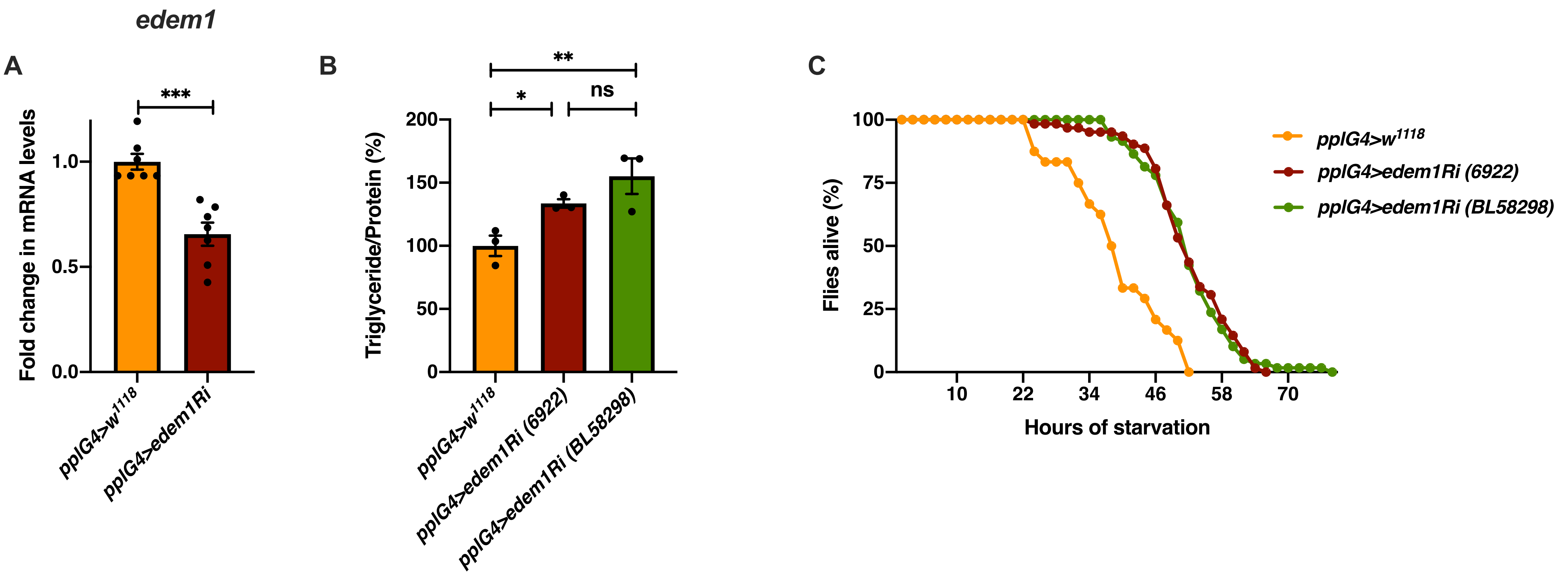


**Fig. S1 Fold change in *edem1* levels in response to *edem1*-RNAi expression in the fat body**

(A) Down-regulation of *edem1* in the fat body led to decreased levels of *edem1* mRNA when compared to control. Data is normalised to *pplGal4>w$^{1118}$* and fold change in *pplGal4>UAS-edem1-RNAi* is shown. [independent biological replicates = 7 P-value between control and *UAS-edem1-RNAi* is 0.0006 (Mann-Whitney test)].

(B) Blocking *edem1* expression using *UAS-edem1-RNAi* (VDRC_6922) and *UAS-edem1-RNAi* (RRID:BDSC_58298) in the fat body led to enhanced triglyceride levels in adult male flies. Data is shown as % ratio of triglyceride to total protein levels, normalised to 100% in *pplGal4>w$^{1118}$* (control) and increase in experimental conditions *pplGal4> UAS-edem1-RNAi* (VDRC_6922) and *UAS-edem1-RNAi* (RRID:BDSC_58298) [independent biological replicates = 3, P-value between control and *UAS-edem1-RNAi* (VDRC_6922) is 0.0410, P-value between control and *UAS-edem1-RNAi* (RRID:BDSC_58298) is 0.0013 and P-value between *UAS-edem1-RNAi* (VDRC_6922) and *UAS-edem1-RNAi* (RRID:BDSC_58298) is 0.3356 (Kruskal-Wallis test followed by Dunn's post-hoc test)].

(C) Enhanced resistance to starvation in adult male flies caused by blocking *edem1* using *UAS-edem1-RNAi* (VDRC_6922) and *UAS-edem1-RNAi* (RRID:BDSC_58298) expression in the fat body. Data shown as percentage of flies of *pplGal4>w$^{1118}$* (control) and *UAS-edem1-RNAi* (VDRC_6922) and *UAS-edem1-RNAi* (RRID:BDSC_58298) which were alive at various time points of starvation [independent biological replicates = 3, number of flies used for control is 24, for *pplGal4>UAS-edem1-RNAi* (6922) is 62 and for *pplGal4> UAS-edem1-RNAi* (BL58298) is 59. P-value between control and *UAS-edem1-RNAi* (VDRC_6922) is <0.001, P-value between control and *UAS-edem1-RNAi* (RRID:BDSC_58298) is <0.001 and P-value between *UAS-edem1-RNAi* (VDRC_6922) and *UAS-edem1-RNAi* (RRID:BDSC_58298) is 0.8256 (Log-rank test), Wald test = 14.11 on df = 1, p<0.001 (cox-proportional hazard analysis)].

**[*P-value *<0.05; ** <0.01,*** <0.001; Data information: In (A-B) data are presented as mean ± SEM*].**

#### Fig. S2

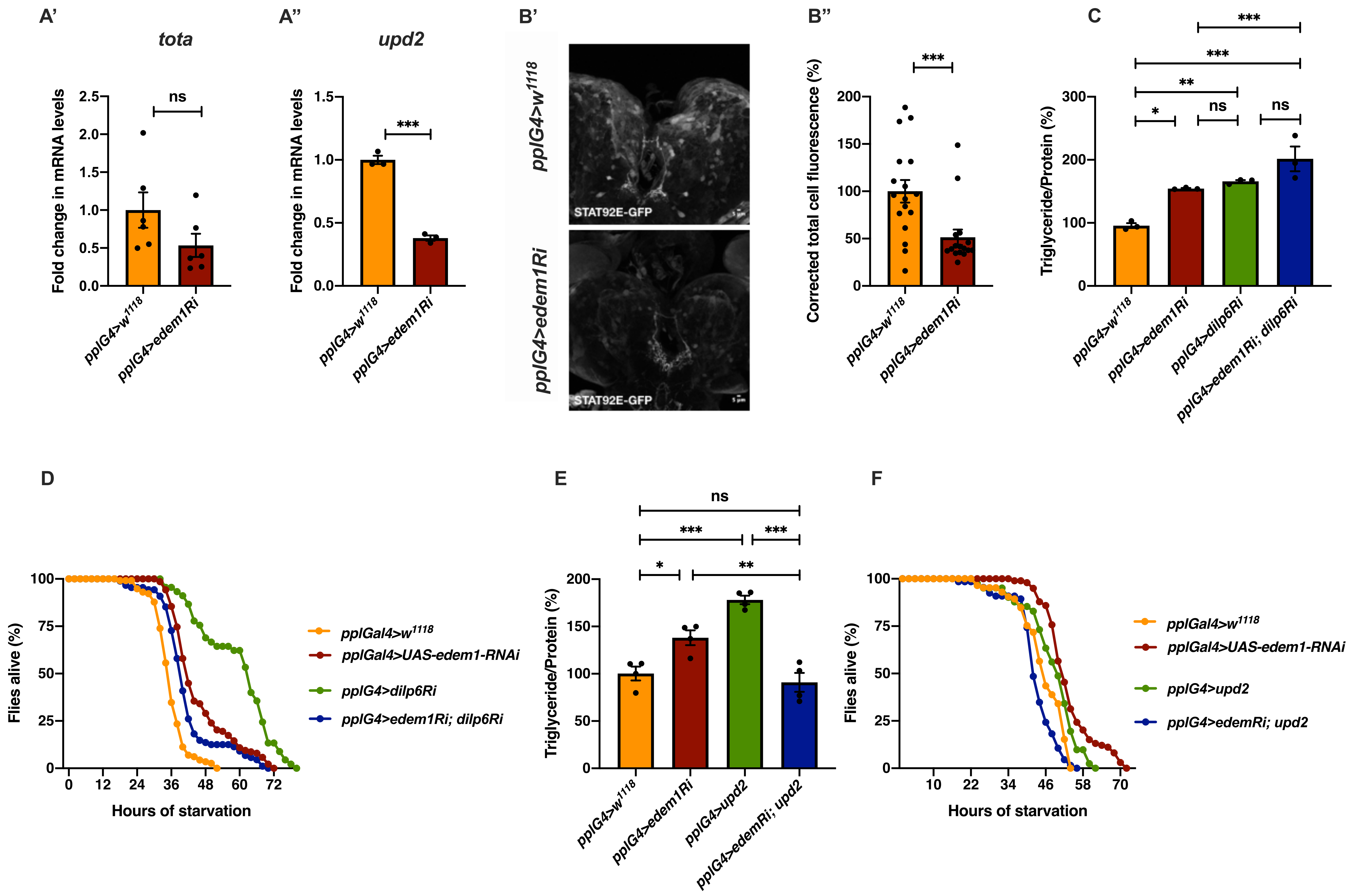

A'
tota
Fold change in mRNA levels
ns
pplG4>w1118
pplG4>edem1Ri
A''
upd2
***
B'
pplG4>w1118
pplG4>edem1Ri
STAT92E-GFP
B''
Corrected total cell fluorescence (%)
***
C
Triglyceride/Protein (%)
pplG4>dilp6Ri
pplG4>edem1Ri; dilp6Ri
D
Flies alive (%)
Hours of starvation
pplGal4>w1118
pplGal4>UAS-edem1-RNAi
pplG4>dilp6Ri
pplG4>edem1Ri; dilp6Ri
E
pplG4>upd2
pplG4>edemRi; upd2
F
pplG4>upd2
pplG4>edemRi; upd2

**Fig. S2 Co-expression of *upd2* but not *dilp6-RNAi* partially rescued the metabolic phenotypes.**

(A) Down-regulation of *edem1* in the larval fat body led to decrease in *totA* mRNA levels (Fig. EV3A'). Data is shown as fold change in mRNA levels, values are normalised to *pplGal4>*$w^{1118}$ and fold change in *pplGal4>UAS-edem1-RNAi* is shown. [independent biological replicates = 6. P-value between control and *UAS-edem1-RNAi* is 0.0649 (Mann-Whitney test)]. Down-regulation of *edem1* in the larval fat body led to decrease in *upd2* mRNA levels (Fig. EV3A''). Data is shown as fold change in mRNA levels, values are normalised to *pplGal4>*$w^{1118}$ and fold change in *pplGal4>UAS-edem1-RNAi* is shown. [independent biological replicates = 6. P-value between control and *UAS-edem1-RNAi* is 0.0003 (Welch's t test)].

(B) Down-regulation of *edem1* in the larval fat body led to decrease in STAT92E-GFP expression (B'). Shown are the representative images of anti-GFP antibody staining in larval brains of *pplGal4>*$w^{1118}$ [independent biological replicates = 17] and *pplGal4>UAS-edem1-RNAi* [independent biological replicates = 16]. (B'') Corrected total cell fluorescence values are normalised to *pplGal4>*$w^{1118}$ and fold change in *pplGal4>UAS-edem1-RNAi* is shown. [P-value between control and *UAS-edem1-RNAi* is 0.0017 (Mann-Whitney test)].

(C) Co-expression of *UAS-dilp6-RNAi* in the fat body did not rescue enhanced stored fat levels caused by *edem1*-RNAi. Data is shown as % ratio of triglyceride to total protein levels, values are normalised to *pplGal4>*$w^{1118}$ and fold change in *pplGal4>UAS-edem1-RNAi, pplGal4> UAS-dilp6-RNAi* and *pplGal4> UAS-edem1-RNAi; UAS-dilp6-RNAi* is shown. [independent biological replicates = 3, P-value between control and *UAS-edem1-RNAi* is 0.0061, P-value between control and *UAS-dilp6-RNAi* is0.0013**,** P-value between *UAS-edem1-RNAi* and *UAS-edem1-RNAi*, *UAS-dilp6-RNAi* is 0.0019**,** P-value between *UAS-edem1-RNAi* and *UAS-dilp6-RNAi* is >0.9999**,** P-value between *UAS-dilp6-RNAi* and *UAS-edem1-RNAi*, *UAS-dilp6-RNAi* is >0.9999 and P-value between control and *UAS-edem1-RNAi*, *UAS-dilp6-RNAi* is <0.001 (Kruskal-Wallis test followed by Dunn's post-hoc test)].

(D) Co-expression of *UAS-dilp6-RNAi* in the fat body did not rescue increased starvation resistance caused by *edem1*-RNAi. Data is shown as percentage of flies which were alive at various time points of starvation in the following genotypes - *pplGal4>*$w^{1118}$, *pplGal4>UAS-edem1-RNAi, pplGal4>UAS-dilp6-RNAi* and *pplGal4> UAS-edem1-RNAi; UAS-dilp6-RNAi.* [independent biological replicates = 3, number of flies used for control is 115, for *pplGal4>UAS-edem1-RNAi* is 138, for *pplGal4>UAS-dilp6-RNAi* is 45 and for *pplGal4> UAS-edem1-RNAi*, *UAS-dilp6-RNAi* is 88. P-value between control and *UAS-edem1-RNAi* is <0.001, P-value between control and *UAS-dilp6-RNAi* is <0.001**,** P-value between *UAS-edem1-RNAi* and *UAS-edem1-RNAi*, *UAS-dilp6-RNAi* is 0.0035**,** P-value between *UAS-edem1-RNAi* and *UAS-dilp6-RNAi* is <0.001**,** P-value between *UAS-dilp6-RNAi* and *UAS-edem1-RNAi*, *UAS-dilp6-RNAi* is <0.001 and P-value between control and *UAS-edem1-RNAi*, *UAS-dilp6-RNAi* is <0.001 (Log-rank test), Wald test = 33.26 on df = 1, p<0.001 (cox-proportional hazard analysis)].

(E) Overexpression of *upd2* in the fat body rescued enhanced stored fat levels caused by *edem1*-RNAi. Data is shown as % ratio of triglyceride to total protein levels, values are normalised to *pplGal4>*$w^{1118}$ and fold change in *pplGal4>UAS-edem1-RNAi*, *pplGal4>UAS-upd2-EGFP* and *pplGal4> UAS-edem1-RNAi; UAS-upd2-EGFP* is shown. [independent biological replicates = 4, P-value between control and *UAS-edem1-RNAi* is 0.001, P-value between control and *UAS-upd2-EGFP* is <0.001**,** P-value between *UAS-edem1-RNAi* and *UAS-edem1-RNAi*, *UAS-upd2-EGFP* is 0.0002**,** P-value between *UAS-edem1-RNAi* and *UAS-upd2-EGFP* is 0.0370**,** P-value between *UAS-upd2-EGFP* and *UAS-edem1-RNAi*, *UAS-upd2-EGFP* is <0.001 and P-value between control and *UAS-edem1-RNAi*, *UAS-upd2-EGFP* is >0.9999 (Kruskal-Wallis test followed by Dunn's post-hoc test)].

(F) Overexpression of *upd2* in the fat body rescued increased starvation resistance caused by *edem1*-RNAi. Data is shown as percentage of flies which were alive at various time points of starvation in the following genotypes - *pplGal4>*$w^{1118}$, *pplGal4>UAS-edem1-RNAi, pplGal4>UAS-upd2-EGFP* and *pplGal4> UAS-edem1-RNAi; UAS-upd2-EGFP.* [independent biological replicates = 3,number of flies used for control is 85, for *pplGal4>UAS-edem1-RNAi* is 99, for *pplGal4> UAS-upd2-EGFP* is 41 and for *pplGal4> UAS-edem1-RNAi*, *UAS-upd2-EGFP* is 66. P-value between control and *UAS-edem1-RNAi* is <0.001, P-value between control and *UAS-upd2-EGFP* is <0.001**,** P-value between *UAS-edem1-RNAi* and *UAS-edem1-RNAi*, *UAS-upd2-EGFP* is <0.001**,** P-value between *UAS-edem1-RNAi* and *UAS-upd2-EGFP* is 0.0073**,** P-value between *UAS-upd2-EGFP* and *UAS-edem1-RNAi*, *UAS-upd2-EGFP* is <0.001 and P-value between control and *UAS-edem1-RNAi*, *UAS-upd2-EGFP* is 0.003 (Log-rank test), Wald test = 9.01 on df = 1, p=0.003 (cox-proportional hazard analysis)].

**[*P-value *<0.05; ** <0.01,*** <0.001; Data information: In (A, B''-C and E) data are presented as mean ± SEM*].**

#### Fig. S3

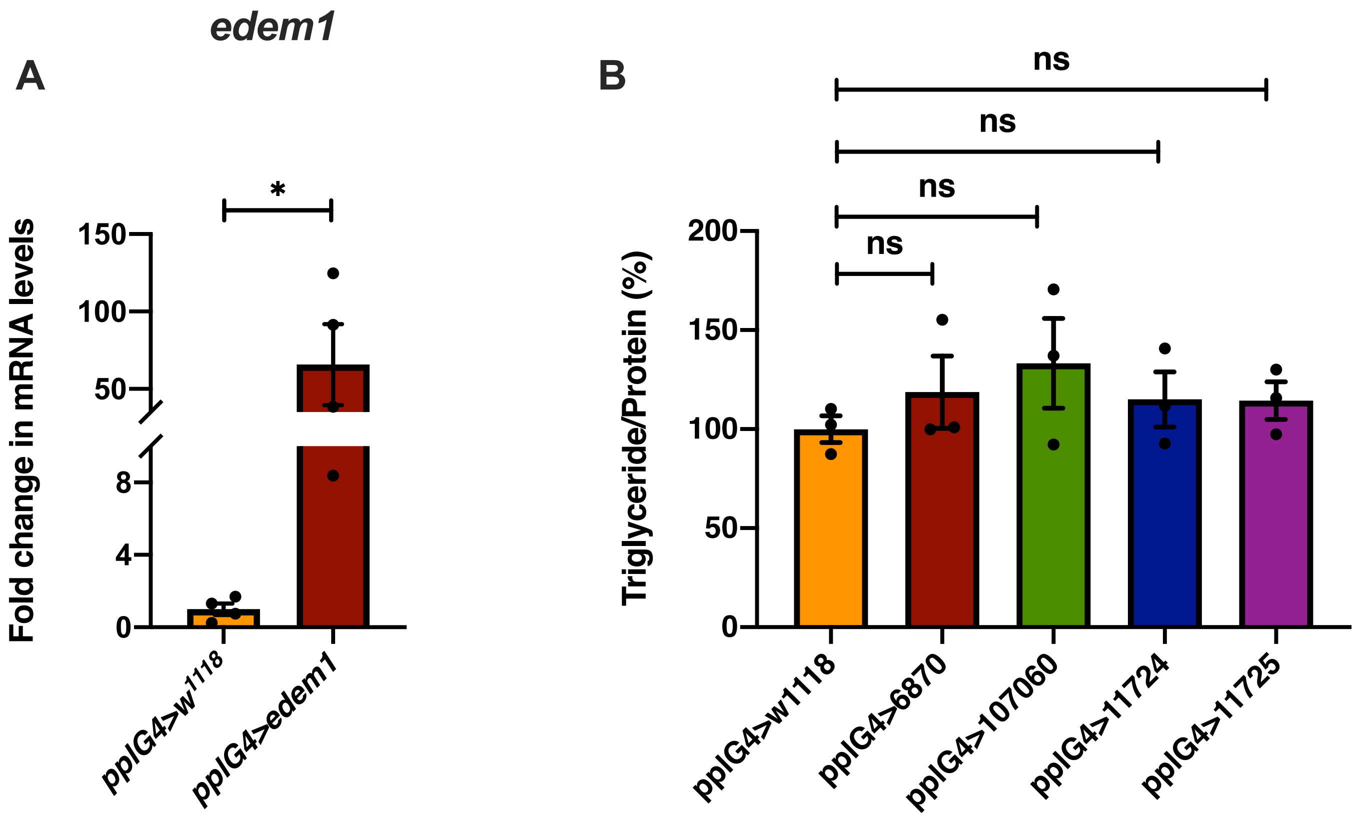


**Fig. S3 Fold change in *edem1* mRNA levels in response to *edem1* over expression in the fat body**

(A) Over-expression of *edem1* in the fat body led to increased levels of *edem1* mRNA when compared to control. Data is normalised to *pplGal4>w$^{1118}$* and fold change in *pplGal4>UAS-edem1-RNAi* is shown. [independent biological replicates = 4 P-value between control and *UAS-edem1-RNAi* is 0.0477 (Unpaired t test)].

(B) Blocking ERAD using RNAi against sip3 and herp in the fat body did not affect the triglyceride levels 5-day old flies. Data is shown as % ratio of triglyceride to total protein levels, normalised to 100% in *pplGal4>w$^{1118}$* (control) and experimental conditions *pplGal4>UAS-sip3-RNAi* (*6870* and *107060)* and *pplGal4>UAS-herp-RNAi* (*11724* and *11725)* [independent biological replicates = 3, P-value between control and *pplG4>6870, pplG4>107060, pplG4>11724* and *pplG4>11725* is >0.9999 (Kruskal-Wallis test followed by Dunn's post-hoc test)].

**[*P-value *<0.05; ** <0.01,*** <0.001; Data information: In (A-B) data are presented as mean ± SEM*].**
